# Building Goal-Directed Cognitive Graphs

**DOI:** 10.64898/2026.02.18.706628

**Authors:** Adithya Gungi, Pradyumna Sepúlveda, Ines F. Aitsahalia, Marta Blanco-Pozo, Kiyohito Iigaya

**Affiliations:** Department of Physics, Columbia University, New York, NY; Department of Psychiatry, Columbia University Irving Medical Center, New York, NY; The Italian Academy for Advanced Studies, Columbia University, New York, NY; Center for Theoretical Neuroscience and Zuckerman Institute, Columbia University, New York, NY; 2CNC Program, Stanford University, Stanford, CA; James H. Clark Center for Biomedical Engineering & Sciences, Stanford University, Stanford, CA; New York State Psychiatric Institute, New York, NY; Data Science Institute, Columbia University, New York, NY

## Abstract

Flexible behavior requires building internal structured models from experience to support goal-directed action. Although environmental transition statistics accumulate gradually, how they are used to construct a compact goal-directed graph remains unclear. We introduce the Sparse Cognitive Graph (SCG), a reinforcement-learning framework that separates gradual transition learning from the sparse directed graph that governs valuation and action selection. In the SCG, transition statistics accumulate in a dense predictive representation, while nonlinear selection determines which transitions are expressed as graph edges. Consequently, gradual strengthening can trigger discrete reorganization of graph topology and abrupt behavioral shifts. Across human reward and transition revaluation tasks, the SCG explains bimodal and trimodal behavioral regimes as emergent consequences of distinct graph configurations. Across human and mouse two-step tasks, dynamic graph reconfiguration captures canonical reward-by-transition interactions without requiring mixtures of control systems. In mice, transitions preceding reward strengthened more rapidly, biasing graph topology toward reward-directed paths. Temporally precise optogenetic dopamine stimulation produced behavioral effects consistent with accelerated graph edge formation predicted by the SCG. The model further generates a testable prediction: graph topology determines the geometry of low-dimensional population activity. Directed acyclic graphs yield activity concentrated at graph entry and goal states, whereas cyclic graphs produce periodic, grid-like structure. Together, these findings identify reward-dependent graph reorganization as a computational principle that reconciles stable predictive learning with efficient goal-directed control.

## Introduction

A hallmark of biological intelligence is the ability to extract relational structure from experience and use that structure to guide flexible, goal-directed behavior. Converging evidence suggests that such structure can be formalized as graph-like internal models (“cognitive graphs”),^1–9^ in which states are linked by directed relationships specifying routes to goals, such as reward outcomes. Neural signatures of predictive and relational coding are distributed across hippocampal^10–14^ and prefrontal circuits,^15–22^ across spatial and abstract domains.

Across these systems, a subtle, yet important contrast emerges. Hippocampal–entorhinal circuits exhibit dense transition representations consistent with successor-like coding,^10–14^ whereas prefrontal populations often express a more sparse, task-relevant structure during planning and choice.^15–22^ Although hippocampal output contributes to prefrontal representations,^23–25^ it remains unclear how dense transition representations are transformed to the compact, goal-directed graph that appears to guide behavior.

Successor representations (SR) provide a principled account of how predictive relationships are acquired, specifying how states accumulate discounted expectations of future occupancy.^3,26,27^ In standard formalism, valuation and choice are computed directly from this dense transition representation that updates gradually with experience. However, behavioral evidence suggests that organisms do not always act on fully expressed predictive maps. Humans and other animals often rely on compressed, task-relevant structure.^28–31^ Moreover, behavior frequently exhibits abrupt, regime-like shifts, even when experience accumulates gradually.^32–34^ These observations suggest a computational dissociation: predictive transition statistics may strengthen gradually, while the internal graph guiding behavior reorganizes discretely. This dissociation raises a central computational question: what determines which transitions are incorporated into the behaviorally expressed graph, and which remain latent within the predictive representation?

Reward and dopamine are strong candidates for modulating this transformation. Dopaminergic signals encode reward prediction errors,^35–37^ unexpected transitions and latent states,^38–41^ regulate effective learning rates,^42,43^ and shape plasticity across hippocampal–prefrontal circuits.^44–47^ Yet, how reward and dopamine shape goal-directed structure remains unsolved.

Here we introduce the Sparse Cognitive Graph (SCG), a reinforcement-learning framework that formalizes this transformation. In the SCG, transition statistics accumulate continuously in a dense transition representation, while behavior is governed by a sparsified directed graph derived through nonlinear selection. This separation allows gradual transition learning to yield discrete reorganization of graph topology and abrupt behavioral shifts. We show that this mechanism explains multimodal individual differences in human reward and transition revaluation,^48^ reproduces canonical two-step task signatures in humans and mice^49,50^ without invoking arbitration between distinct controllers, and provides a computational interpretation of dopaminergic manipulation as modulation of graph construction. The framework further generates testable predictions linking graph topology to low-dimensional population structure.

Together, these results identify reward-dependent graph reorganization as a computational mechanism that reconciles gradual predictive learning with efficient goal-directed behavior by transforming graded transition statistics into discrete, behaviorally expressed graph structure.

## Results

### Building sparse cognitive graphs through reinforcement learning

We formalize the Sparse Cognitive Graph (SCG), a reinforcement-learning framework in which gradual transition learning and nonlinear sparse graph construction are computationally separable. Gradual transition learning refers to the accumulation of transition statistics in a dense representation, whereas nonlinear sparse graph construction refers to the selective formation of a sparse directed graph that governs valuation and choice.

As an agent interacts with an environment composed of directed state transitions (**Fig. 1A**), it incrementally updates a dense *transition representation W* (**Fig. 1B**). This transition representation (TR) captures how each state predicts its successor states under temporal discounting, and is learned via a temporal-difference rule closely related to successor-representation (SR) learning.^3,26^

**Figure 1.**
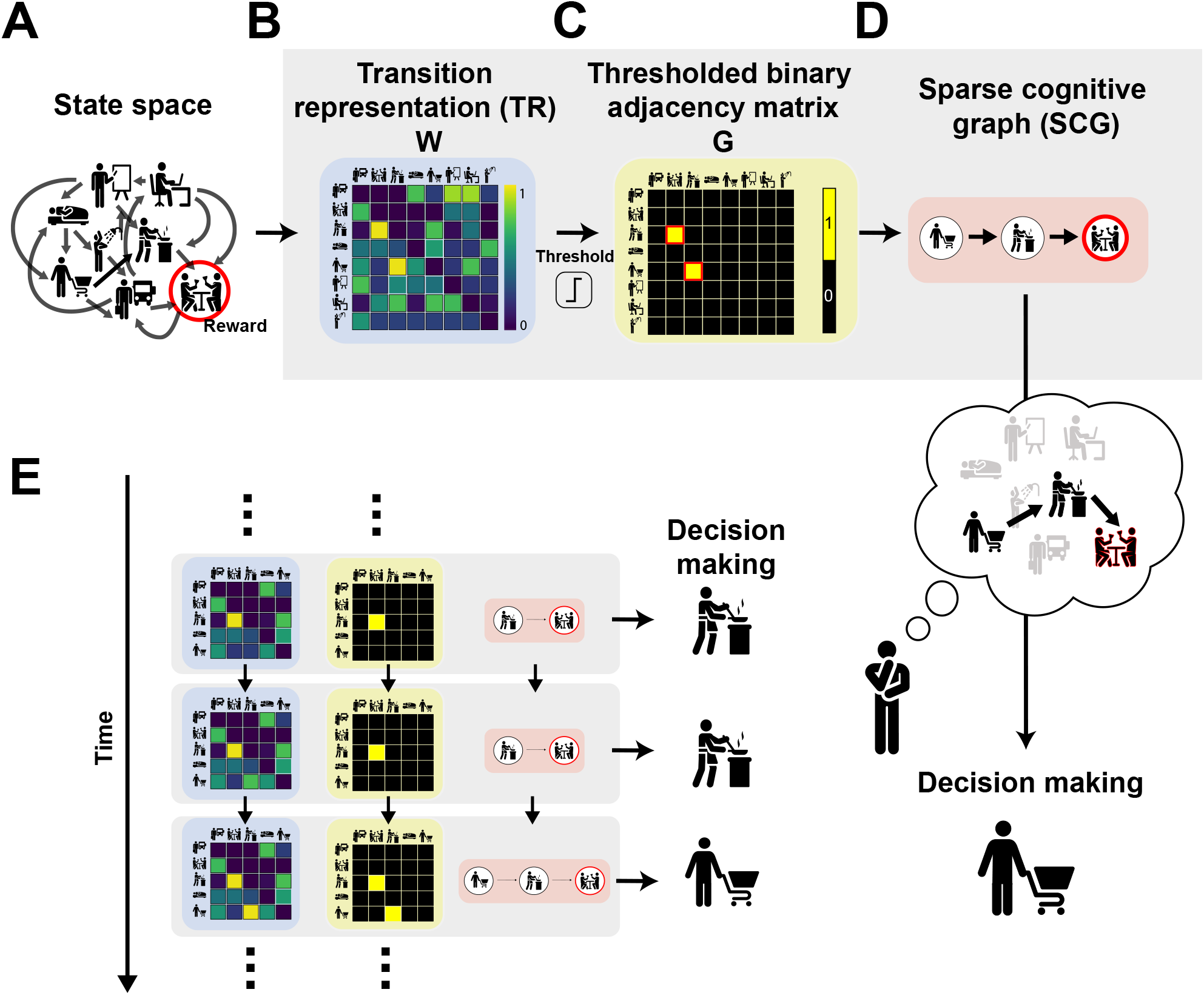
Sparse Cognitive Graph (SCG): separating gradual transition learning from graph-based action selection. (**A**) An agent experiences sequential transitions between latent states. (**B**) Gradual transition learning: the agent updates a dense transition representation (TR) *W* that integrates discounted successor predictions over transition experience. In the formulation studied here, this TR *W* does not itself govern valuation or action selection. (**C**) Nonlinear graph construction: after each update, TR *W* is transformed by a nonlinear selection rule (threshold ζ) to produce a sparse directed adjacency matrix *G*. (**D**) The adjacency matrix *G* defines the Sparse Cognitive Graph. This graph governs valuation and action selection. (**E**) Throughout learning, the dense transition representation *W* and the sparse graph *G* are updated in parallel. TR *W* continuously integrates new experience, whereas *G* can reorganize discretely as transitions in *W* cross threshold.

We adopt the TR rather than the classic SR because our goal is to construct an explicit directed graph of experienced transitions. Whereas the SR encodes discounted future state occupancy, the TR is anchored to experienced one-step transitions and therefore provides a direct substrate for graph construction. This distinction is clearest in the limit of zero discounting: the TR converges to the empirical one-step transition matrix, whereas the SR converges to the identity matrix (see Methods). As a result, the TR naturally supports extraction of directed relational structure from ongoing experience.

After each experienced transition, the updated TR *W* is mapped to a sparse binary adjacency matrix *G* by a nonlinear selection rule implemented here as thresholding at ζ (**Fig. 1C**). Entries of *W* that exceed threshold are incorporated as edges (*G*_*ij*_ = 1), whereas all others are suppressed (*G*_*ij*_ = 0). In graph-theoretic terms, *G* defines a directed graph whose nodes correspond to states and whose edges denote reliably predicted transitions (**Fig. 1D**). We refer to this adjacency matrix *G* as the Sparse Cognitive Graph (SCG), a computational abstraction of the structure hypothesized to guide behavior. In the present work, we analyze a minimal instantiation using a fixed threshold applied uniformly across transitions and trials.

This nonlinear selection step yields a compact internal structure that can support efficient planning and is consistent with capacity limits on maintaining task structure through working memory.^51^ Importantly, thresholding should be understood as an algorithmic abstraction of nonlinear selection rather than a literal hard cutoff, and could arise from multiple biological mechanisms (online or offline), including competitive gating, synaptic stabilization, or replay-mediated pruning. Here we adopt the simplest formulation while remaining agnostic about neural implementation.

The model maintains both the dense TR *W* and the sparse graph *G*, which evolve jointly throughout learning (**Fig. 1E**). Each experienced transition triggers (i) a temporal-difference update to *W* (gradual transition learning) and (ii) an online structural update to *G*. This co-evolution preserves long-range predictive information in *W* while allowing graph *G* to reorganize discretely as transition strengths cross threshold. Because our focus is on how inferred graph structure governs behavior, we assume that valuation and action selection operate on the graph *G*, rather than directly on the dense representation *W*. This contrasts with standard SR-based approaches, in which behavior is typically derived directly from dense predictive representations.^26^

Motivated by evidence that reward modulates learning rates,^50,52–57^ we allow the TR learning rate to depend on whether the observed transition is followed by reward, *α*_→*R*_, or not, *α*_→*NoR*_. Importantly, we impose no a priori ordering between these rates; instead, we test for such asymmetries empirically.^50^ If *α*_→*R*_ *> α*_→*NoR*_, transitions preceding reward accumulate predictive strength more rapidly in *W*, increasing their likelihood of being incorporated into *G*. Through this mechanism, reward biases graph construction toward pathways leading to valuable outcomes, without explicitly encoding goals.

### Nonlinear Graph Construction Generates Discrete Behavioral Regimes

To test whether nonlinear sparse graph construction can account for discrete patterns of human structure learning, we applied the SCG to reward and transition revaluation data originally reported by Momennejad and colleagues.^48^ These tasks probe how individuals update relational structure under limited experience. Participants exhibit strikingly discrete response patterns, an empirical feature that has lacked a mechanistic explanation. We therefore asked whether such patterns could arise from nonlinear graph construction in the SCG.

In both tasks, participants first learned two three-step sequences:

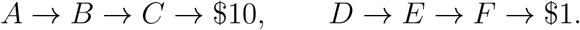

In the reward revaluation task (**Fig. 2A**), terminal rewards were swapped during revaluation without revisiting the initial states:

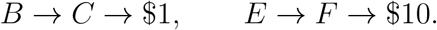

**Figure 2.**
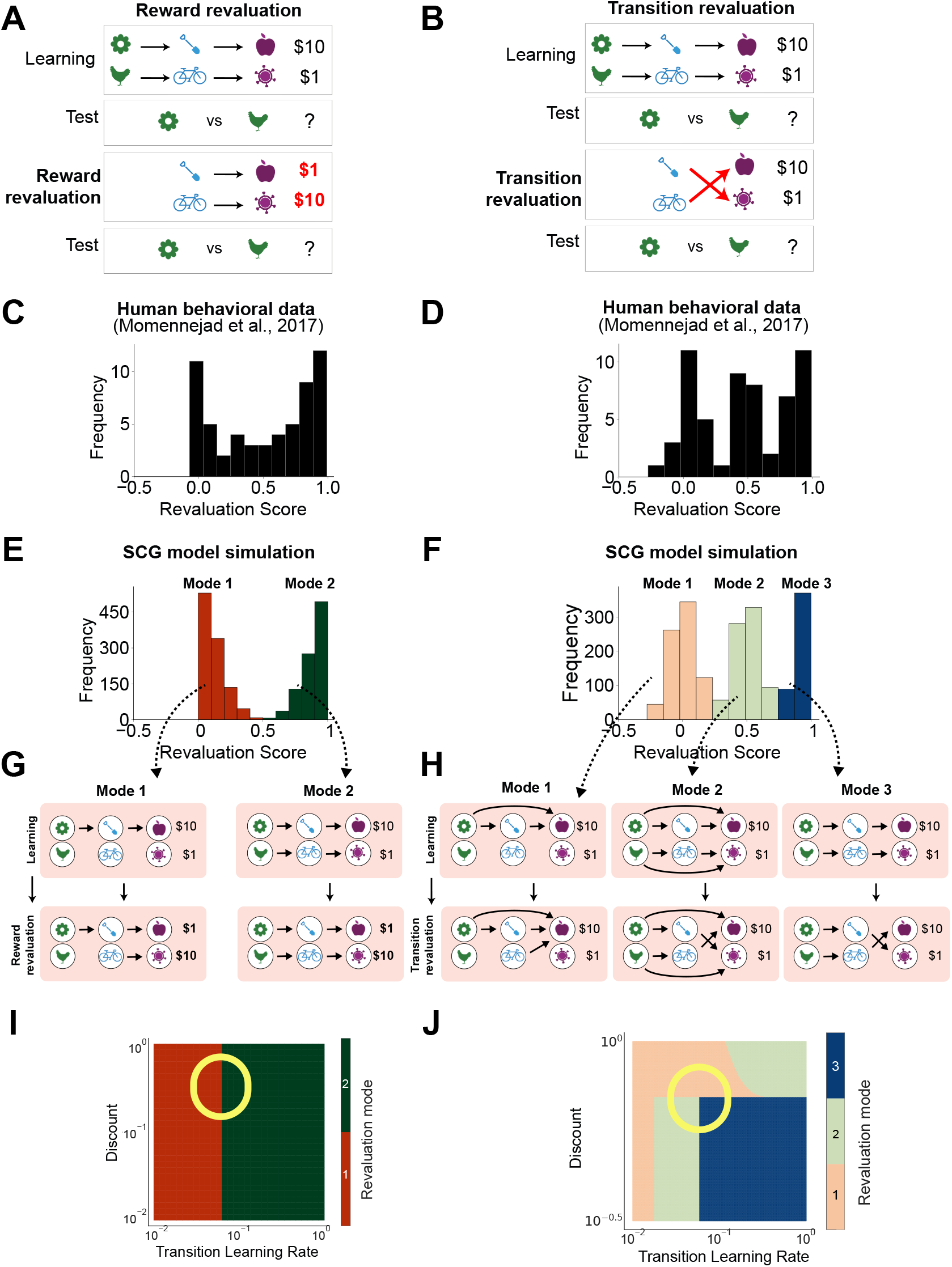
Sparse cognitive graphs produce discrete behavioral modes in human revaluation tasks.^48^. (**A**) Reward revaluation task. Participants learned two independent three-step sequences leading to high ($10) or baseline ($1) reward. During revaluation, terminal rewards were swapped without re-exposure to starting states. (**B**) Transition revaluation task. The intermediate transition structure was altered while terminal rewards remained fixed. (**C**) Reward revaluation data showing a bimodal distribution of revaluation scores. (**D**) Transition revaluation data showing a trimodal distribution. (**E**,**F**) SCG simulations reproduce these distributions despite unimodally distributed model parameters. (**G**) Graph topologies corresponding to the two behavioral modes in reward revaluation (Mode 1 and Mode 2). (**H**) Graph topologies corresponding to the three behavioral modes in transition revaluation (Mode 1–3). (**I**,**J**) Phase diagrams demonstrating discrete regimes of graph topology emerging as model parameters vary smoothly. Colors in (E) correspond to those in (I), and colors in (F) correspond to those in (J), indicating matched behavioral modes. Circles indicate parameters used for simulations.

Participants therefore had to update starting-state values by propagating new reward information through previously learned transition structure.

In the transition revaluation task (**Fig. 2B**), intermediate transitions were altered while rewards remained fixed:

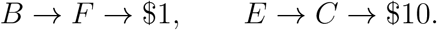

Again, starting states were not re-experienced, requiring inference over latent structural change rather than direct reinforcement.

Behavioral responses exhibited striking multimodality.^48^ Revaluation was quantified using a normalized revaluation score reflecting change in relative preference between *A* and *D*. In reward revaluation, scores clustered near 0 (no change) and 1 (complete reversal) (**Fig. 2C**). In transition revaluation, scores clustered near 0, 0.5 (indifference), and 1 (**Fig. 2D**). These discrete behavioral modes suggest that participants may differ not only quantitatively, but qualitatively, in the internal structure they express.

We asked whether such discrete modes could emerge from nonlinear sparse graph construction in the SCG. In simulations, model agents were drawn from a unimodal parameter distribution and exposed to identical task experiences. Despite this, the SCG produced discrete graph configurations *G* (**Fig. 2G,H, Fig. S1**), which in turn generated bimodal and trimodal behavioral distributions (**Fig. 2E,F**). Thus, multimodal behavior arose not from multimodal parameters, but from discrete graph topologies induced by nonlinear graph construction.

Phase diagram analyses clarified this mechanism (**Fig. 2I,J**). Smooth variation of model parameters revealed discrete regions of graph topology. Because sparse graph construction involves a nonlinear thresholding operation, small parameter changes near regime boundaries induced abrupt reorganization of the graph *G*, and therefore abrupt shifts in choice. This nonlinearity allows multimodal behavioral distributions to emerge even when underlying parameters are unimodally distributed.

In these simulations, transition learning rates were allowed to differ between transitions yielding $10 and $1 outcomes, with the smaller outcome treated as baseline. Learning-rate modulation therefore reflects the presence (*α*_→*R*_) versus absence (*α*_→*NoR*_) of additional reinforcement rather than absolute reward magnitude.

By contrast, standard computational models, including classic SR, model-free temporal difference learning, fully model-based reinforcement learning, and standard mixture models, did not reproduce these discrete behavioral modes under unimodal parameter distributions (**Fig. S2**). Models that rely solely on dense transition representations without nonlinear sparse graph construction produced smoothly varying behavior. These comparisons highlight the necessity of nonlinear graph construction for generating discrete patterns of human structure learning.

Together, these results demonstrate that multimodal behavioral modes can emerge from the nonlinear mapping between gradual transition learning and sparse graph construction, supporting this separation as the core computational principle of the SCG.

### Dynamic Graph Reorganization Generates Reward-by-Transition Choice Patterns

We next asked whether the SCG can account for behavior in the widely studied two-step task.^49^ In this paradigm (**Fig. 3A**), participants choose between two first-stage options, each leading probabilistically to one of two second-stage states via common (70%) or rare (30%) transitions. Second-stage reward probabilities drift slowly over time.

**Figure 3.**
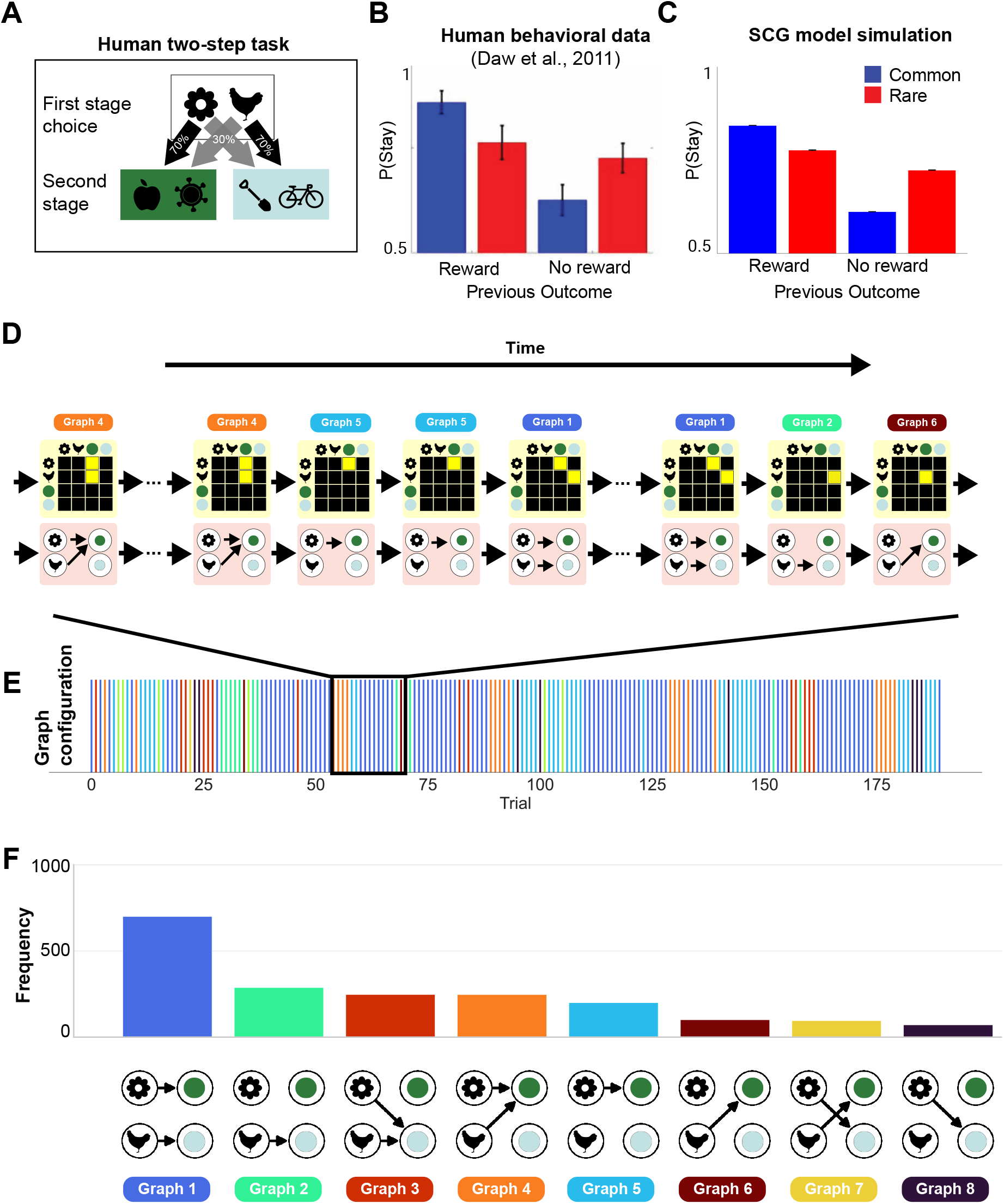
Dynamic graph reorganization generates reward-by-transition interactions in the two-step task. Two-step task.^49^ First-stage choices lead probabilistically to second-stage states (common 70%, rare 30%). Second-stage reward probabilities drift slowly across trials. (**B**) Human stay probability as a function of reward outcome and transition type.^49^ Stay probability reflects the likelihood of repeating the previous first-stage choice. Error bars indicate s.e.m. across participants. Reproduced from.^49^ (**C**) SCG simulations reproduce the canonical reward-by-transition interaction. Error bars indicate s.e.m. across simulations. (**D**) Trial-by-trial reorganization of the expressed graph *G* during simulation. Recent transitions and rewards reshape the graph topology across trials. (**E**) Temporal evolution of graph configurations across trials. Colors denote distinct graph configurations expressed at different points in learning. The corresponding graphs are shown in D and F. (**F**) Distribution of graph configurations across trials in simulation. The most frequent graph (Graph 1) retained only common transitions.

Behavior in this task is often summarized using the stay-probability analysis (**Fig. 3B**), which quantifies the probability of repeating a first-stage choice as a function of the previous trial’s reward outcome and transition type. Model-free learning predicts repetition following reward regardless of transition, whereas model-based learning predicts a reward-by-transition interaction.^49^ Empirical data typically exhibit such an interaction, motivating mixture-based interpretations. However, accumulating evidence indicates that similar behavioral signatures can arise from alternative computational assumptions.^58–61^

We found that the SCG reproduces the canonical reward-by-transition interaction without requiring explicit arbitration between distinct controllers (**Fig. 3C**). In the SCG, behavior is governed by the currently expressed graph *G*, rather than by a fixed transition model or a weighted mixture of value systems. As transition statistics in *W* are updated gradually, nonlinear graph construction produces trial-by-trial reorganization of *G* (**Fig. 3D,E**). Across a session, agents express a distribution of graph configurations (**Fig. 3F**), reflecting dynamic changes in inferred relational structure.

Reconfiguration of the expressed graph within the SCG is sufficient to generate the reward-by-transition interaction observed in human behavior. Crucially, the interaction emerges from the nonlinear mapping between gradual transition learning and sparse graph construction, rather than from combining separate model-based and model-free controllers.

Although alternative accounts have been proposed,^58–60^ these results demonstrate that canonical two-step signatures can arise from dynamic graph inference alone. The SCG thus explains two-step behavior as a consequence of ongoing structural reorganization during learning.

To test this structural interpretation more directly, we next applied the SCG to mouse two-step behavior and evaluated the model quantitatively.

### Reward and Dopamine Bias Graph Construction

To test whether the SCG generalizes across species and to assess its quantitative validity, we applied the model to a mouse two-step dataset with dopaminergic manipulation.^50^ In this task (**Fig. 4A**), mice chose between first-stage options that led probabilistically to second-stage states, with reward probabilities determined by unsignaled block structure.

**Figure 4.**
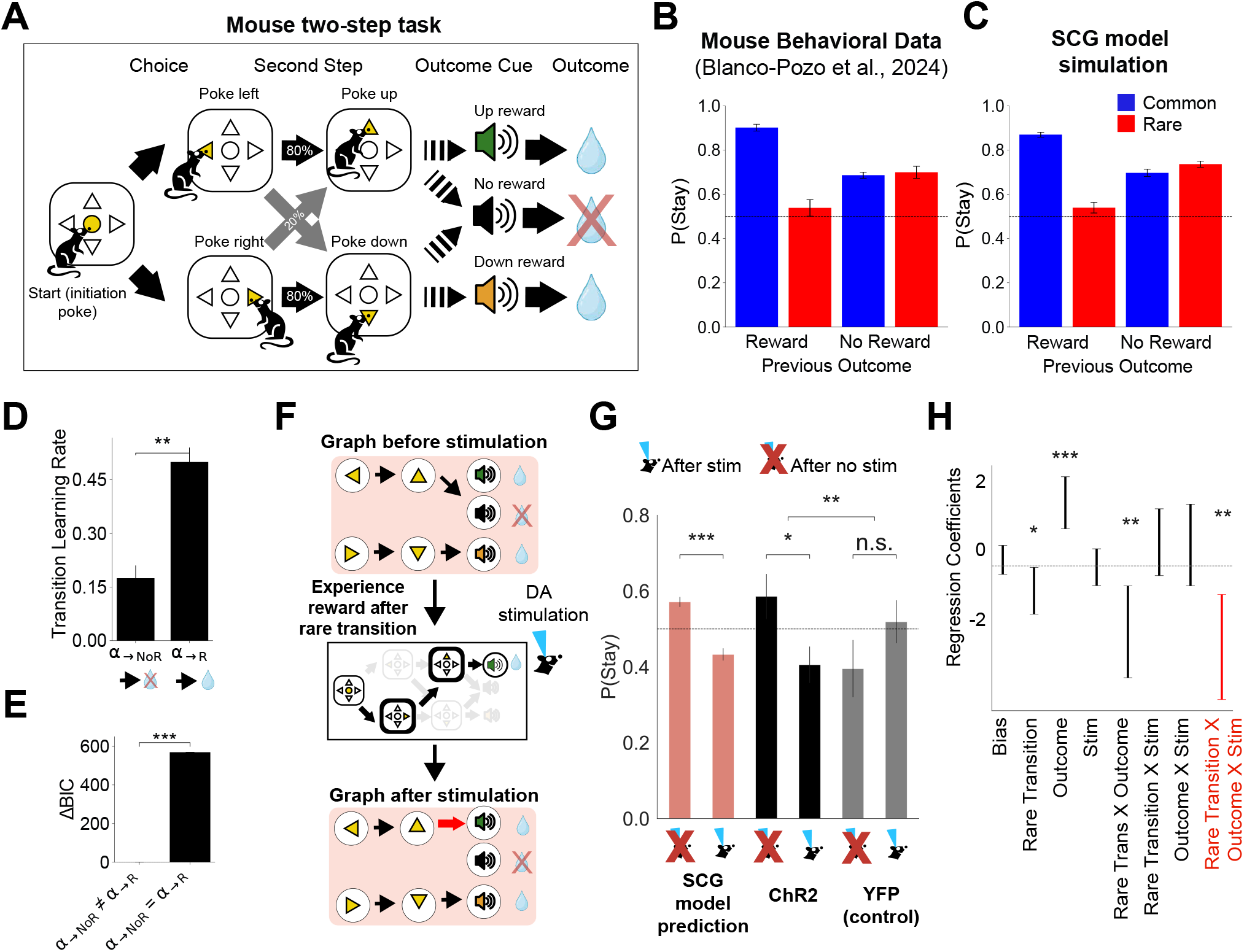
Reward and dopamine bias nonlinear graph construction in the mouse two-step task. Data originally reported in.^50^ (**A**) Mouse two-step task. First-stage choices led probabilistically to second-stage states (common 80%, rare 20%). Reward probabilities followed an unsignaled block structure. (**B**) Mouse choice patterns differed from those observed in humans and were not well explained by standard model-based or model-free reinforcement learning accounts, as previously reported.^50^ (**C**) SCG simulations reproduced the observed mouse behavioral pattern. Model parameters were fit to trial-by-trial choice data.^50^ Error bars indicate s.e.m. across subjects. (**D**) Fitted transition learning rates. Transition learning was stronger following reward than no reward (*α*_→*R*_ *> α*_→*NoR*_). Error bars indicate *±* s.e.m. across subjects. (**E**) Model comparison (iBIC) favored asymmetric transition learning. Smaller values indicate better fits. Error bars indicate cross-sample mean ± s.e.m. (4,000 bootstrap samples from the posterior distribution). See **Fig. S5** for full model comparisons across other candidate models (including the SR and the previously winning model). (**F**) Illustrative example of how optogenetic stimulation of dopamine neurons can induce graph reconfiguration, leading to shifts in subsequent choice. Optogenetic stimulation at the outcome was modeled as an increase in the transition learning rate, analogous to reward. (**G**) Model prediction and behavioral data. The model predicts reduced stay probability following optogenetic stimulation on rewarded rare-transition trials. Behavioral data from the Channelrhodopsin-2 (ChR2) group showed this effect (*p* < 0.05, permutation test), whereas the yellow fluorescent protein (YFP) control group showed no significant effect. The effect was significantly stronger in the ChR2 group than in the YFP control group (*p* < 0.01, permutation test). Error bars indicate s.e.m. across subjects. (**H**) Logistic regression confirming a significant Rare × Reward × Stimulation interaction in ChR2 animals. This supports increased switching behavior following rewarded rare transitions under stimulation. The effect was absent in the control group (see **Fig. S7**). Error bars indicate *±*2 SE.

The original study reported three notable findings.^50^ First, mouse behavior deviated from predictions of standard mixtures of model-based and model-free reinforcement learning (**Fig. 4B**). Second, learning appeared asymmetric following reward versus no reward. Third, optogenetic stimulation of ventral tegmental area dopamine neurons at outcome did not produce behavioral shifts predicted by standard value-update interpretations.

We instantiated the SCG with a state space aligned to the task structure and fit model parameters to trial-by-trial choices using a hierarchical framework.^28,62^ The SCG reproduced the observed behavioral pattern (**Fig. 4C**) through dynamic nonlinear graph construction of graph configurations across trials (**Fig. S3**).

Crucially, fitted parameters revealed that transition learning in *W* was significantly stronger following rewarded transitions than unrewarded transitions (*α*_→*R*_ *> α*_→*NoR*_; permutation test *p* < 0.001; **Fig. 4D**). Model comparison further supported asymmetric transition learning (**Fig. 4E**).

Within the SCG framework, this asymmetry has a specific structural interpretation: reward acceler-ates transition learning for recently experienced transitions, increasing their likelihood of being incorporated as edges in the graph *G*. Thus, reward biases nonlinear graph construction toward pathways that precede valuable outcomes.

By contrast, other standard computational models, including the classic SR, model-free temporal difference learning, and model-based reinforcement learning, were unable to reproduce the observed mouse behavioral patterns (**Fig. S4**), consistent with previous reports.^50^ Formal hierarchical model comparison further favored the SCG over these alternatives, including the best-performing model from the original analysis (a model-based agent with asymmetric reward learning rates;^50^ **Fig. S5**). Notably, the SR model, which relies on dense transition representations without nonlinear graph construction, did not account for the data, highlighting the computational role of nonlinear sparse graph construction.

### Dopamine Stimulation Promotes Graph Reorganization

The original study reported that dopamine stimulation at outcome did not produce the behavioral shifts expected if dopamine acted solely as a reward prediction error signal.^50^ Within the SCG, however, reward has two computational roles: (i) value updating, and (ii) modulation of gradual transition learning, thereby influencing graph construction.

We therefore asked whether modeling dopamine stimulation as a transient increase in transition learning rate would yield a distinct behavioral signature. In the SCG, a single rewarded rare transition is typically insufficient to reorganize the graph *G*. However, if paired with stimulation that amplifies transition learning, the same experience can increase in strength within *W*, exceed threshold, and be incorporated as a new edge in *G*.

This mechanism generates a specific prediction: stimulation following rewarded rare transitions should increase subsequent switching by inducing reorganization of graph topology (**Fig. 4F**). Consistent with this prediction, mice expressing channel-rhodopsin (ChR2) exhibited a reduced stay probability following stimulation on rewarded rare-transition trials, unlike control mice (**Fig. 4G**, **Fig. S6**; *p* < 0.01, permutation test). Regression analyses confirmed a significant Rare × Reward × Stimulation interaction in ChR2 animals (**Fig. 4H**), which was absent in control animals (**Fig. S7**).

Together, these findings support a computational interpretation in which reward and dopamine bias graph construction by modulating transition learning. This mechanism complements established roles of dopamine in value updating while identifying a distinct structural pathway through which dopamine can reorganize internal graph topology and reshape behavior.

### Graph Topology Predicts Low-Dimensional Population Geometry

Beyond behavior, the SCG makes predictions about topology-dependent low-dimensional population signatures arising from graph structure. Previous work has shown that spectral decompositions of dense transition representations, including the SR, can yield grid-like periodic population signatures in spatial environments.^3,63^ These periodic signatures arise when predictive structure approximates translational symmetry.

We therefore asked how population signatures change when predictive information is sparsified into a directed graph. To address this, we analyzed the leading eigenvectors of the expressed adjacency matrix *G*, which reveal dominant low-dimensional population signatures implied by graph topology.

In sparse directed graphs learned in the two-step task (**Fig. 5A–C**), the dominant population signatures were not periodic. Instead, one signature was localized near entry states of directed subgraphs (**Fig. 5D**), while a complementary signature was localized near terminal (goal) states (**Fig. 5E**). These structured signatures reflect directional reachability in the graph: activity concentrates at sources and sinks of the learned topology. We refer to these boundary- and goal-localized representations as *flag-like* signatures to distinguish them from periodic grid-like signatures.

**Figure 5.**
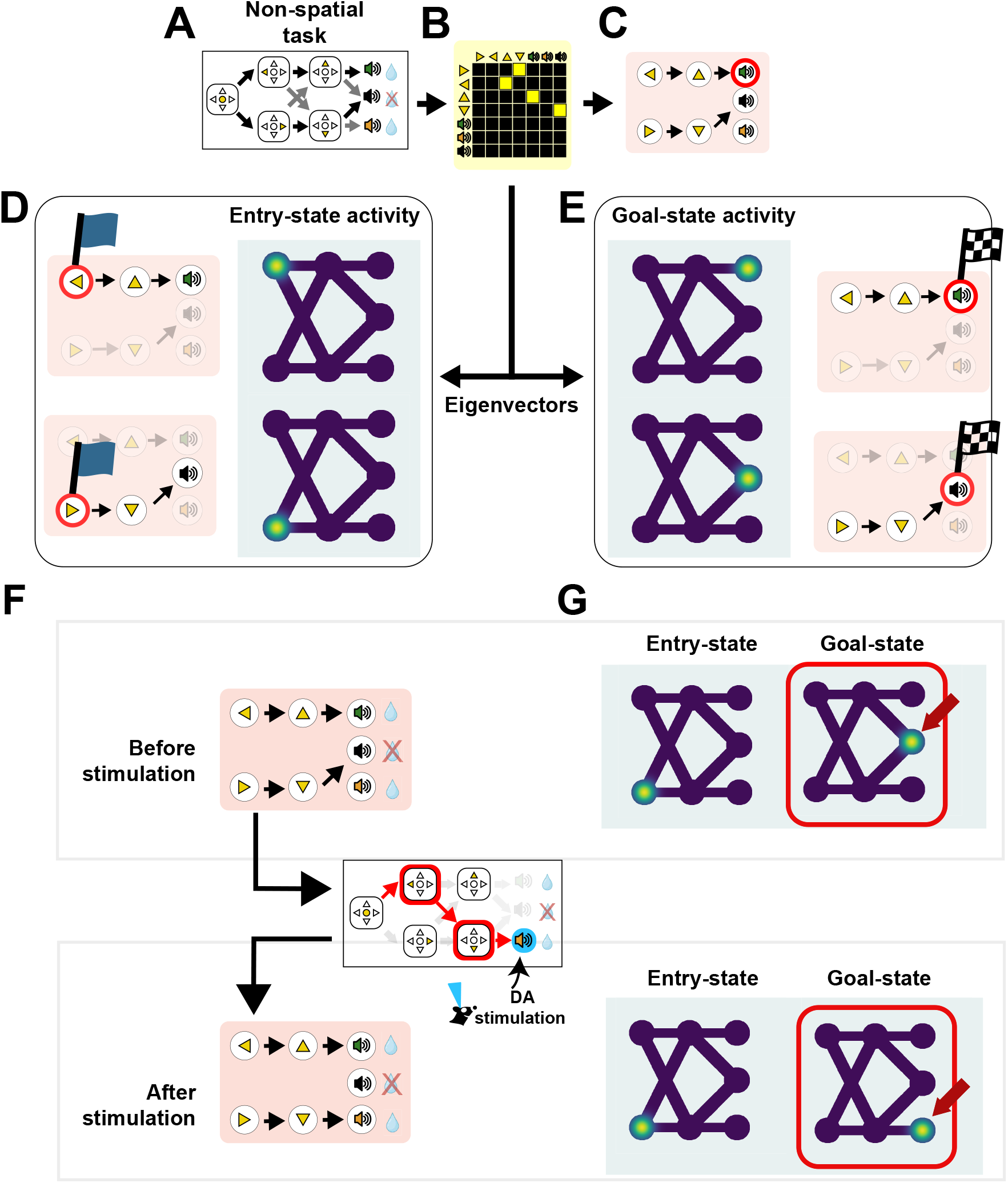
Graph topology predicts low-dimensional population geometry. (**A**) Example two-step task structure. (**B**) Sparse adjacency matrix *G* inferred by the SCG. (**C**) Directed graph corresponding to *G*. (**D**) Predicted low-dimensional population signature localized at graph entry states, arising from right eigenvectors of *G*. Two distinct signatures correspond to the two entry states of separate subgraphs. (**E**) Predicted complementary low-dimensional population signature localized at graph goal states, arising from left eigenvectors of *G*. Two distinct signatures correspond to the two goal states of separate subgraphs. (**F**) Predicted effect of dopamine-driven increases in transition strength: reorganization of graph topology through enhanced edge expression in *G*. (**G**) Predicted consequence of graph reorganization: corresponding changes in low-dimensional population signatures. In this example, goal-state signatures shift in accordance with changes in graph configuration.

Importantly, these predictions concern population-level structure rather than single-neuron tuning. The SCG does not assume neurons explicitly encode “start” or “goal.” Rather, low-dimensional population signatures emerge as a mathematical consequence of graph topology.

Whether periodic (grid-like) or boundary-localized (flag-like) signatures dominate depends on the cyclicity and symmetry of the graph. Directed acyclic graphs emphasize sources and sinks, producing localized signatures. In contrast, when repeated task structure induces a cyclic graph approximating translational symmetry, the SCG predicts periodic low-dimensional signatures even in non-spatial tasks (**Fig. S8A**). Extending the model to two-dimensional navigation produces a similar pattern (**Fig. S8B**). When sparsification is sufficiently weak to preserve approximate symmetry, the SCG reproduces grid-like structure analogous to that derived from the SR.^3^

Low-dimensional structure can also arise from the dense transition representation *W*. However, because *W* integrates graded transition strength across indirect paths, its dominant modes reflect smooth predictive geometry. In contrast, sparsification into *G* imposes directed reachability constraints, accentuating asymmetries between sources and sinks and amplifying boundary- and goal-localized structure. Thus, the qualitative regime shift from periodic to boundary-localized signatures can arise from nonlinear graph construction rather than from transition learning alone.

Thus, the SCG unifies boundary-localized and periodic population signatures within a single computational framework. Dense, symmetric transition structure gives rise to periodic signatures, whereas sparse, directed graph construction emphasizes goal-relevant boundaries. In this way, sparsification transforms predictive geometry into task-aligned population structure.

### Dopamine-Dependent Graph Reorganization Predicts Changes in Population Geometry

Building on the relationship between graph topology and population structure, the SCG makes a concrete prediction: dopamine-dependent learning should reorganize low-dimensional population signatures by altering the topology of the graph *G*.

In the SCG, dopamine modulates gradual transition learning in the TR *W*. When predictive strength for a specific transition is amplified, such as through optogenetic stimulation paired with reward, that transition becomes more likely to exceed threshold and be structurally expressed as an edge in *G*. Because low-dimensional population signatures are determined by the topology of *G*, any dopamine-driven change in expressed edges necessarily alters the associated population signatures.

We illustrate this prediction in the two-step task (**Fig. 5F,G**). Dopamine-dependent amplification of transition learning alters the strength of specific edges in *W*, increasing their likelihood of inclusion in *G*. Such edge changes reorganize the topology of the expressed graph, even when underlying experience differs only minimally.

These topological shifts, in turn, reorganize the leading low-dimensional population signatures derived from *G*. In directed graphs, these signatures reflect the distribution of sources and sinks within the structure. Modifying a single directed edge can alter reachability relationships, thereby reshaping the boundary- and goal-localized population signatures predicted at the systems level.

This mechanism yields a testable prediction: pairing dopamine stimulation with specific transitions should produce coordinated changes in graph topology, low-dimensional population activity patterns, and subsequent choice behavior. Importantly, this prediction does not require dopamine to directly encode goals or value. Rather, dopamine modulates transition learning, which in turn reshapes graph topology and the geometry of internal representations.

Thus, the SCG provides a mechanistic link between dopamine-dependent modulation of transition learning and coordinated changes in population structure and behavior within a unified computational framework.

## Discussion

### From Gradual Transition Learning to Goal-Directed Structure

Flexible behavior requires extracting predictive regularities from experience and organizing them into compact internal models that guide choice. A central unresolved question is how dense transition representations, consistent with successor-like coding in hippocampal circuits,^3,27^ are transformed into the compact directed relational structure expressed during decision making.^15,17,18^ Existing reinforcement learning frameworks explain how predictive statistics are accumulated and how reward shapes value estimates, but they do not specify how gradual transition learning is converted into the structured graph that governs behavior.

The Sparse Cognitive Graph provides a computational framework for understanding this transformation. In the SCG, transition statistics accumulate gradually in a dense representation *W*, whereas valuation and action selection operate on a sparsified directed graph *G* constructed through nonlinear selection. This separation implies a simple principle. Transition representations in *W* may evolve gradually, yet behavior can reorganize abruptly when the topology of the graph *G* changes. This dissociation allows predictive knowledge to accumulate stably over time, while enabling rapid structural reconfiguration and computationally efficient planning and decision making by restricting valuation to a sparsified graph.

### Nonlinear Sparse Graph Construction as a Mechanism of Discrete Behavioral Mode Shifts

Abrupt or step-like changes in behavior are common and have motivated proposals that learning occurs in discrete or all-or-none events.^32–34^ The SCG provides a computational reconciliation of these observations. Transition learning in *W* proceeds incrementally, yet behavior is governed by the sparsified graph *G*, constructed through nonlinear selection. Because this mapping from *W* to *G* is nonlinear, small quantitative changes in accumulated predictive strength can produce abrupt reorganization of graph topology.

Discrete behavioral modes therefore emerge from structural reconfiguration rather than representational jumps. As observed in human revaluation tasks,^48^ multimodal patterns of choice can arise even when underlying learning parameters vary smoothly. The essential computation is the separation between gradual predictive learning and nonlinear graph construction at the level of decision making.

### Neural and Behavioral Dissociations

A direct consequence of this separation is that neural signatures of gradual transition learning need not align temporally with behavioral change. Transition statistics encoded in *W* may strengthen gradually and be detectable in neural population activity consistent with successor-like coding in hippocampal circuits,^3^ even when behavior remains unchanged. Behavioral reorganization occurs only when accumulated transition strength crosses the threshold and reorganizes the topology of the graph *G*.

This framework therefore predicts systematic dissociations between neural evidence of learning and observable behavior. Neural signatures of statistical transition learning may precede behavioral change, lag behind it, or transiently diverge from it, depending on when thresholds are crossed and graph topology reorganizes. Such discrepancies have been observed in learning paradigms where neural indicators of associative learning emerge gradually before behavior changes appear in a step-like manner (e.g.,^57^). The SCG provides a computational account of these phenomena without requiring arbitration between multiple learning systems or an explicit latent state inference.

### Reward and Dopamine Bias Graph Construction

Our analyses suggest that reward and dopamine influence behavior not only through value updating, but by biasing which predictive transitions become incorporated into the graph that governs choice. In the SCG, asymmetric transition learning rates are sufficient to shift which transitions are retained in *G*, thereby altering graph topology. This interpretation is consistent with a broad literature implicating dopamine in modulating effective learning rates.^42,43^ In the mouse two-step dataset,^50^ fitted parameters supported stronger transition learning following reward than no reward, and model comparison favored this asymmetry. Modeling optogenetic dopamine stimulation as a transient amplification of transition learning generated a distinct behavioral prediction that was confirmed empirically.

Within this framework, dopamine need not explicitly encode graph structure or goals. Instead, reward-dependent modulation of transition learning alters the probability that specific transitions are incorporated into the graph. Through this mechanism, dopaminergic modulation of transition learning can reorganize the expressed graph in the SCG and reshape behavior. This account complements established theories of reward prediction errors^35–37^ and aligns with evidence that dopamine signals unexpected transitions or state prediction errors,^38–41^ while identifying a structural pathway through which dopamine influences decision-making.

### Retrospective Reorganization Without Backward Inference

Reward-triggered reorganization of the graph *G* in the SCG can produce behavioral dynamics that resemble retrospective causal inference. Models such as ANCCR propose that reward receipt triggers restructuring of inferred causal relationships.^57^ In the SCG, however, transition learning remains prospective and incremental. What changes is which transitions become incorporated into the graph governing behavior. Reward-dependent modulation of transition learning increases the likelihood that recently experienced transitions are retained in *G*. Once incorporated, these transitions alter the relational topology of the graph and reshape subsequent evaluation. The SCG can therefore generate apparent retrospective reorganization without requiring explicit backward inference.

### Relation to Generative Relational Frameworks

The SCG complements generative relational frameworks such as the Tolman–Eichenbaum Machine^4^ and Clone Structured Cognitive Graph,^5,64^ which emphasize inference over latent relational structure in hippocampal circuits. These approaches focus on how relational maps are constructed, generalized, and aligned across contexts. By contrast, the SCG formalizes how reward-dependent transition learning determines which learned relationships are incorporated into the compact directed graph that governs choice. Rather than modeling generative inference over latent structure, the SCG specifies a reinforcement-constrained mechanism through which predictive knowledge becomes behaviorally operative.

### Representational Decoupling and Computational Advantage

Sparsification has computational implications. As shown by formal complexity analysis (please see Methods), dense predictive matrices scale quadratically with the size of the state space, whereas sparse directed graphs scale with the number of retained edges.^65,66^ By restricting planning to the graph *G*, the SCG provides a route from predictive richness to computational efficiency.

Because valuation and policy evaluation operate on the sparsified graph *G* rather than the full predictive matrix *W*, the effective dimensionality of decision computation is reduced to the number of retained edges. Planning complexity therefore scales with task-relevant structure instead of the combinatorial size of the state space. This separation permits dense predictive learning to proceed without imposing corresponding computational costs at the level of action selection, allowing predictive representations to remain high-dimensional while keeping goal-directed planning computationally tractable.

This perspective aligns with evidence that humans rely on simplified relational structure during planning,^28,29^ adopt resource-rational approximations,^67–69^ and treat highly probable transitions as effectively deterministic.^30,31^ Within this framework, sparsification is not merely compression, but a structural operation that shapes qualitative features of behavior by determining which relationships are retained in the graph that governs choice.

### Hippocampal–Prefrontal Interactions and Graph Construction

The SCG is a computational-level framework and does not assume that neural circuits explicitly encode adjacency matrices. Rather, it proposes that goal-directed behavior reflects nonlinear selection of predictive structure functionally equivalent to graph construction. Within this perspective, the model offers a principled framework for interpreting hippocampal–prefrontal interactions.

One possibility consistent with existing evidence is that hippocampal circuits maintain relatively dense predictive representations resembling successor maps^3^ (corresponding to *W*), while cortical or frontostriatal circuits express more compact, task-constrained relational structure (corresponding to *G*) that shapes prediction and action. Such a division of labor would naturally implement the computational advantages described above, allowing hippocampus to preserve high-dimensional predictive structure while prefrontal circuits operate on a sparsified graph that supports efficient planning and flexible reconfiguration. Although the precise circuit-level transformation remains to be established,^21^ the SCG provides a concrete framework for investigating how predictive representations in hippocampus may be transformed into compact relational structure in prefrontal cortex to guide goal-directed behavior.

### Topology-Dependent Population Geometry

Within the SCG, graph topology predicts the geometry of low-dimensional population structure. In directed acyclic graphs of the type studied here, right eigenvectors emphasize entry-point (source) states, whereas left eigenvectors emphasize terminal (goal) states. Cyclic graphs with approximate translational symmetry give rise to periodic grid-like modes.^3,63^ These signatures arise from directed reachability constraints and therefore constitute testable predictions about population-level organization rather than assumptions about single-neuron tuning.

Experimentally, the model predicts that tasks with acyclic, goal-directed structure should elicit boundary- and goal-localized low-dimensional modes in regions expressing compact relational structure, such as prefrontal and frontostriatal circuits, whereas cyclic or translationally symmetric tasks should produce periodic grid-like modes. Dopamine-dependent modulation of transition learning is further predicted to reorganize this geometry: manipulations that selectively strengthen specific transitions should induce measurable shifts in dominant population modes.

Boundary- and context-related signals are widely reported in hippocampal–subicular circuits,^70–72^ and suppressing hippocampal–subicular output to prefrontal cortex accelerates the emergence of generalized schema representations.^25^ Within the SCG framework, reducing strong context anchoring would facilitate formation of more abstract, transferable cognitive graph structure across tasks. Conversely, the goal-localized signatures predicted by the SCG conceptually overlap with outcome- and goal-related signals observed in prefrontal and frontostriatal systems,^73^ suggesting a potential correspondence between graph-defined terminal states and neural representations involved in goal evaluation.

### Limitations and Future Directions

The present work analyzes a minimal instantiation of graph construction using a fixed nonlinear selection rule and assumes that valuation operates exclusively on the graph *G*. In biological systems, graph construction may depend on context, uncertainty, or neuromodulatory state, and behavior may integrate information from both dense transition representations and sparsified graphs. Extending the framework to incorporate adaptive graph construction or dynamic interaction between these representational regimes is an important direction for future work. Dopaminergic systems operate across multiple timescales.^74,75^ Incorporating multi-scale discounting into transition learning could support hierarchical graph organization, linking distributional coding to structured internal models. Although our analyses focused on reward-based learning, the same principle naturally extends to aversive domains, where dopamine also plays a role.^76^ Threat-related modulation of transition learning may similarly accelerate graph formation, potentially accounting for rapid, few-shot structural reorganization observed in aversive contexts.^77^ Finally, direct neural tests of the theory will require simultaneous measurement of gradual transition learning and expressed relational structure to determine whether dopaminergic modulation reorganizes low-dimensional population signatures as predicted. Identifying neural correlates of both dense predictive representations and sparsified graph structure within the same task will be critical for adjudicating between representational and structural accounts of learning.

In sum, by separating gradual transition learning from nonlinear graph construction, the Sparse Cognitive Graph identifies a computational mechanism through which dense predictive knowledge is reorganized into compact goal-directed structure. This framework accounts for multimodal human behavior, cross-species decision signatures, dopamine-dependent structural reorganization, and neural–behavioral dissociations within a unified model, while generating falsifiable predictions about topology-dependent low-dimensional population signatures.

## Methods

### M.1 Computational Model of Sparse Cognitive Graph (SCG)

#### Model overview

Our framework formalizes how an agent constructs a Sparse Cognitive Graph (SCG) from experience using a reinforcement-learning algorithm that separates gradual transition learning from nonlinear graph construction.

The model maintains two complementary internal representations throughout learning: (1) a dense predictive *transition representation* (TR), denoted *W*, which summarizes how each state predicts its successors under temporal discounting; and (2) a sparse directed adjacency matrix *G*, obtained by applying a nonlinear graph-construction rule to *W* after each update.

Whereas the transition representation *W* provides a graded record of experienced transition statistics through gradual transition learning, the adjacency matrix *G* defines the Sparse Cognitive Graph that supports planning, valuation, and action selection. Nonlinear graph construction yields a compact directed structure that prioritizes reliably experienced transitions and evolves online as experience accumulates.

In the formulation used here, decisions are based on the graph *G*, rather than directly on the dense transition representation *W*, reflecting the hypothesis that biological agents may rely on simplified internal structure rather than dense predictive matrices.

The Methods are organized as follows. We first describe the SCG learning algorithm independently of specific tasks, followed by task-specific implementations and statistical analyses.

#### Transition representation (TR) *W* and gradual transition learning

For each experienced transition *s*_*t*_ → *s*_*t*+1_, the model performs gradual transition learning in the transition representation using a temporal-difference rule anchored on the observed next state:

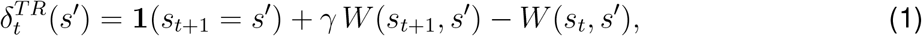

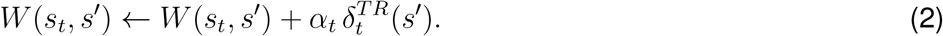

This transition representation tracks experienced transitions while propagating predictive strength forward through the discount factor *γ*. Gradual transition learning therefore integrates transition statistics continuously while maintaining sensitivity to multi-step structure via temporal discounting.

In contrast to the classic successor representation (SR), which tracks discounted future state occupancy, the transition representation *W* remains more localized around experienced one-step transitions and is therefore well suited for graph construction of a sparse directed structure. In the limit *γ* = 0, the update for *W* reduces to the FORWARD algorithm,^78^ corresponding to learning the empirical one-step transition matrix.

#### Reward-modulated gradual transition learning

To allow reward to influence which transitions become expressed in the graph, we permitted learning rates to differ depending on whether a transition was followed by reward:

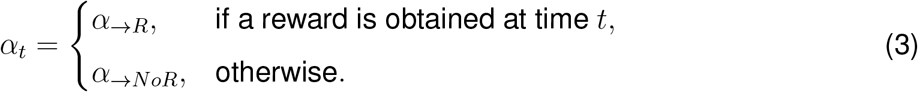

No ordering was imposed between *α*_→*R*_ and *α*_→*NoR*_. If the data support *α*_→*R*_ *> α*_→*NoR*_, then transitions preceding reward accumulate greater predictive strength in *W*, biasing subsequent graph construction toward reward-aligned structure. This asymmetry is therefore an empirical result rather than an a priori assumption.

#### Construction of the Sparse Cognitive Graph

After each update of the transition representation *W*, the agent constructs a sparse directed graph by applying a fixed threshold ζ:

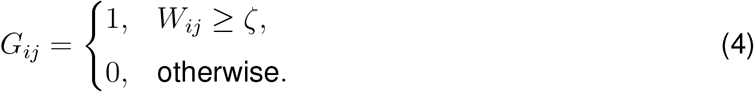

This step retains transitions that have accumulated sufficient predictive strength in *W*. Because graph construction is applied online after each transition, the SCG evolves dynamically as experience is acquired.

The adjacency matrix *G* is then treated as the internal Sparse Cognitive Graph used for valuation and action selection. By contrast, computing decisions directly from *W* yields an SR-like value readout from a dense predictive representation. The graph-construction step therefore enables direct evaluation of the hypothesis that behavior relies on sparse cognitive graphs rather than dense predictive maps.

Accordingly, we explicitly compare SCG agents with standard SR agents to assess the necessity of graph construction across the tasks analyzed.

#### Reward estimation and value computation

Expected reward for each state is learned via a standard delta rule:

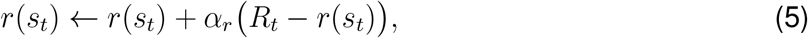

where *R*_*t*_ is the reward observed at time *t* and *α*_*r*_ is the reward-learning rate.

To determine which reward states are reachable from a given state under the current graph, we compute a finite-horizon reachability matrix:^65,66^

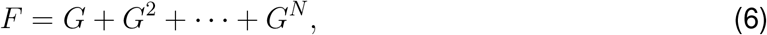

where matrix multiplication and addition are defined over the Boolean semiring, and *N* is chosen to bound or exceed the graph diameter for the task.

In our simulations, where the state space was small, we computed this expression explicitly and binarized the result, setting all nonzero entries to 1. Thus, *F*_*ij*_ = 1 indicates that state *i* can reach state *j* along at least one directed path of length at most *N*.

However, importantly, explicit matrix-power computation like Eq.(6) is not required in general. The same reachability relation can be obtained efficiently using standard graph traversal algorithms,^66^ such as depth-first search, which scales linearly with the number of graph edges rather than quadratically with the number of states.

State values are then computed as

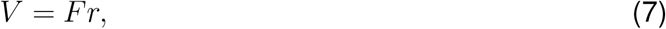

so that *V* (*s*) reflects the aggregate reward reachable from *s* through the SCG.

This value readout is intentionally minimal and serves to demonstrate how the learned SCG can support goal-directed evaluation. Alternative value computations, such as max-based, policy-weighted, or control-dependent formulations, are possible, particularly in graphs with branching structure. Importantly, the central claims of this work do not depend on the specific form of value computation or choice policy. Our contribution concerns how predictive transition structure is accumulated in *W* and nonlinearly expressed as a compact directed SCG, and how reward and dopamine may bias graph formation.

#### Relation to the classic successor representation

The classic successor representation (SR) *M* estimates discounted future state occupancy under the current policy:^26^

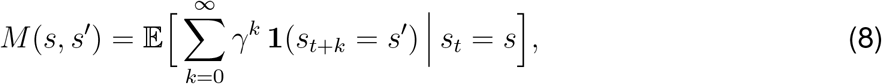

and can be learned incrementally via temporal-difference updates.

In the present work, our objective is to construct a sparse directed graph whose edges correspond directly to learned transitions between states. For this purpose we use the transition representation *W*, which accumulates predictive evidence primarily for observed one-step transitions while still propagating predictive strength via discounting. Entries *W* (*s, s*^*′*^) therefore admit a natural interpretation as directed edges following graph construction.

This distinction is clearest when *γ* = 0. In this limit, *W* reduces to the empirical one-step transition matrix, directly reflecting experienced transitions. By contrast, the SR collapses to the identity matrix, encoding only trivial self-occupancy. Thus, *W* naturally supports graph construction even in regimes where the SR does not provide informative relational structure.

Mathematically, learning *W* and learning the SR *M* are closely related; asymptotically, under this update rule, *M* ≈ *I* + *γW*, where *I* is the identity matrix.

We emphasize that this choice reflects modeling convenience rather than a limitation of the SR framework. More sophisticated graph-extraction procedures could be applied to SR representations. The central hypothesis of the SCG concerns nonlinear graph construction and behavioral use of predictive transitions in a sparse directed graph, rather than the specific predictive operator employed.

#### Computational efficiency

In the SCG, downstream inference and value computation operate on the sparse graph rather than dense predictive matrices. Let *N*_SCG_ denote the number of nodes in the learned SCG and |*E*| the number of directed edges. Storage scales as *O*(|*E*|), with 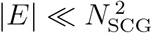 in structured environments. Reachability-based inference can be implemented using standard graph traversal algorithms with computational cost *O*(|*E*| + *N*_SCG_).^65,66^

By contrast, dense transition representations such as *W* or the SR operate over the full state space. Let *N*_SR_ denote the number of states represented in these dense models. Such representations require 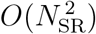 storage and compute values via dense linear operations on the reward vector, incurring 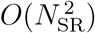 cost per evaluation.^3,26^

Thus, the SCG reduces computational cost through two mechanisms: (i) sparsification of relational structure via graph construction 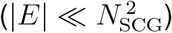, and (ii) compression of the effective state representation (*N*_SCG_ ≪ *N*_SR_).

Importantly, this efficiency does not eliminate the need for a dense predictive representation. Maintaining both *W* and *G* serves distinct computational roles. The dense representation *W* supports gradual temporal-difference updates and integration of multi-step transition statistics across the full state space, retaining predictive information even for transitions not currently expressed in the graph. The sparsified graph *G* is derived from *W* and used for downstream planning and value computation. This separation allows statistical learning to proceed in the high-dimensional space of *W* while restricting decision computation to the compact structure of *G*.

### M.2 Alternative models

To situate SCG within the broader reinforcement-learning literature, we implemented four standard alternatives: (1) the classic successor representation (SR), (2) a purely model-free temporal-difference (TD) learner, (3) a fully model-based (MB) planner, and (4) a hybrid MB/MF mixture. All alternative models were evaluated on the same revaluation and two-step tasks and analyzed using the same revaluation-score and stay-probability measures.

#### classic successor representation (SR)

The successor representation (SR) matrix *M* encodes discounted future state occupancy,

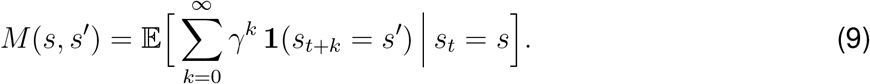

We used a standard one-step temporal-difference update that enforces the SR Bellman equation,

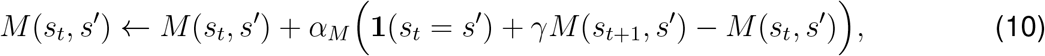

which includes an explicit identity term reflecting current-state occupancy. In parallel, expected rewards were learned via a delta rule,

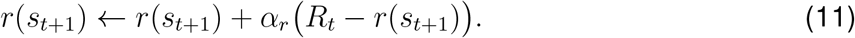

State values were then computed as

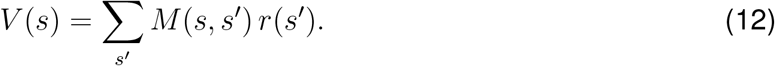

Compared to SCG, the SR encodes a dense predictive map of discounted state occupancy, rather than a sparse directed graph of transitions.

#### Model-free TD learner

The model-free learner updates scalar state values via:

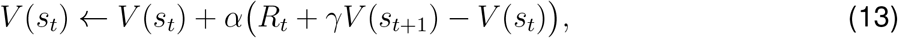

with no explicit transition model. This makes it unable to propagate revaluation through unexperienced starting states.

#### Model-based reinforcement learning

We applied a standard form of MB agent to the two-step task, where asymptotic knowledge of the transition model is given to the agent.^49,50^ To understand the variability of learning, we additionally applied a version of the MB agent that learns the transition model *T* ^78^.^48^ After each observed transition *s*_*t*_ → *s*_*t*+1_, the corresponding entry in *T* is incremented and the row is renormalized to form empirical conditional probabilities.^78^ Given *T* and learned reward estimates, values are obtained by tabular policy evaluation under the learned dynamics.

#### Hybrid MB/MF mixture

In the revaluation experiment, the hybrid agent estimates model-based and model-free values, to estimate the respective ratings (see below), which are combined following the expression :

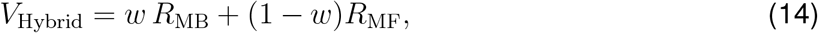

with weighting parameter *w*.

### M.3 Task descriptions

We evaluated the model on previously published behavioral tasks and datasets.

#### Human reward and transition revaluation tasks^48^

We modeled the reward and transition revaluation tasks of Momennejad et al.^48^ using a six-state environment {*A, B, C, D, E, F*}. States *C* and *F* served as terminal states associated with fixed monetary outcomes of 10 (bonus reward) and 1 (baseline outcome), respectively. During initial learning, agents experienced the deterministic structure

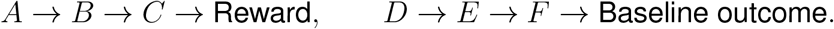

Each sequence was presented 20 times, matching the original experiment. In the experiments, participants reported their relative preference between the two starting states *A* and *D* using a continuous rating scale. These ratings were stored as values that lie between 0 and 1.

In the **reward revaluation** condition, terminal outcomes were swapped while the transition structure was preserved,

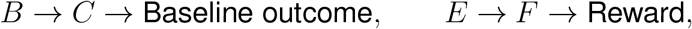

again for 20 presentations; the initial states *A* and *D* were omitted during revaluation.

In the **transition revaluation** condition, the second-step transitions were swapped while terminal outcomes remained unchanged,

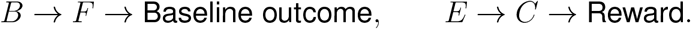

After the revaluation stage, participants were asked to report their relative preference between the two starting states *A* and *D*. The experimental revaluation score was defined as the difference between the post- and pre-relearning preference ratings.^48^

#### Human two-step task.^49^

The standard human two-step task^49^ comprised a first-stage choice between two options, followed by probabilistic transitions to one of two second-stage states. First-stage choices (*A* or *B*) led to second-stage states (*C* or *D*) with common and rare transitions: *P* (*C*|*A*) = 0.7, *P* (*D*|*A*) = 0.3, *P* (*C*|*B*) = 0.3, *P* (*D*|*B*) = 0.7. At the second stage, participants chose between two cues. Reward probabilities for each cue drifted independently over time according to bounded random walks with Gaussian increments (step size *σ* = 0.025) and reflecting boundaries [0.25, 0.75], updated every trial.

#### Mouse two-step task and optogenetic stimulation.^50^

We reanalyzed the publicly available mouse two-step task dataset published in.^50^ Here we summarize the task structure and stimulation protocols relevant for the present analyses. Mice initiated each trial by poking a lit center port and then made a first-stage choice between left and right ports. On 75% of trials both options were available (free-choice), whereas on 25% only a single option was available (forced-choice). Each choice led with 80% (common) vs. 20% (rare) probability to one of two second-step ports (up/down), signaled by distinct auditory and visual cues; the mapping from first-to second-step states was fixed within animals and counterbalanced across animals. Entering the active second-step port triggered a second-step-specific outcome cue indicating reward (either up- or down-reward cue) or no reward (a shared no-reward cue). Reward probabilities at second-step ports varied by block. Biased blocks (80%/20%) ended when an exponentially weighted moving average of correct choices (time constant of 8 free-choice trials) exceeded 75%; balanced blocks (50%/50%) lasted 20–30 trials before pseudo-random switching. Trials ended with a 2–4 s intertrial interval.

Optogenetic stimulation was delivered through implanted optical fibers targeting dopaminergic neurons. In separate experimental sessions, stimulation was applied either at the second-step onset or at the outcome onset. In each session, stimulation occurred on 25% of trials with two constraints: (i) the following trial was always free-choice, and (ii) at least two non-stimulated trials followed each stimulation event. Twelve DAT-Cre mice were tested (5 YFP controls and 7 ChR2 experimental animals). Further procedural details are provided in.^50^

### M.4 Model applications to tasks

#### Human reward and transition revaluation tasks

We applied our model to,^48^ where transition learning rates differed between trials yielding the bonus ($10) and baseline outcomes ($1), with the baseline outcome treated as a reference level such that learning-rate modulation reflected the presence (*α*_→R_) versus absence (*α*_→NoR_) of additional reinforcement.

To characterize qualitative regimes of SCG behavior, we computed phase diagrams over model parameters. To generate **Figure 2IJ**, we varied the baseline transition learning rate *α*_→NoR_ and the discount factor *γ* sampled logarithmically from 10^−2^ to 1, with other parameters fixed at *α*_→R_ = 0.4, *α*_*r*_ = 0.9, and *ζ* = 0.7. For each parameter set, we simulated the full reward or transition revaluation protocol and computed values for the two starting states before revaluation (*pre*) and after revaluation (*post*). Following,^48^ the model computes a latent preference signal for *D* over *A* at time *τ* ∈ {pre, post} as the value difference Δ*V*_*τ*_ = *V*_*τ*_ (*D*) − *V*_*τ*_ (*A*). To mirror the bounded behavioral rating, the reported preference was modeled by mapping this latent signal through a Gaussian cumulative distribution function (probit link), yielding a continuous rating between 0 and 1, following.^48^ The generated ratings, *R*_*τ*_ were used to define the revaluation score within the model as the change in preference,

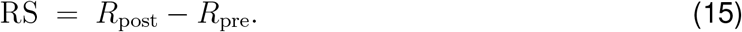

Thus, RS = 0 indicates no change in preference, RS *>* 0 indicates a shift toward *D* (reversal relative to an initial preference).

For the parameter ranges considered here, the model satisfied *V*_pre_(*A*) *> V*_pre_(*D*) at the end of initial learning, consistent with the data in.^48^ We therefore characterized qualitatively distinct revaluation behavioral modes based on the relative ordering of starting-state values after relearning: (i) *V*_post_(*A*) *> V*_post_(*D*), indicating that the initial preference was preserved; (ii) *V*_post_(*A*) < *V*_post_(*D*), indicating a reversal of preference; and (iii) *V*_post_(*A*) ≈ *V*_post_(*D*), indicating indifference between the two starting states.

To generate multimodal preference distributions (**Figure 2EFGH**), we drew parameters from unimodal Gaussian distributions centered near phase boundaries. For reward revaluation, we set *µ*_*α*_ = 0.055 and *µ*_*γ*_ = 0.68, and for transition revaluation, *µ*_*α*_ = 0.042 and *µ*_*γ*_ = 0.68. Variances were set to 0.001 for both parameters, with zero off-diagonal covariance. To mirror the behavioral measure (a noisy preference rating bounded between 0 and 1), following,^48^ we assumed that the revaluation score on each trial was sampled from a Gaussian distribution with mean equal to the model-predicted revaluation score (RS), with noise *σ* = 0.15 for reward revaluation and *σ* = 0.1 for transition revaluation.

We also simulated alternative models to generate the phase diagrams shown in **Fig. S2**. For the mixture of model-free and model-based reinforcement learning, the weighting parameter *w* ∈ [0, 1] was drawn from a Gaussian distribution 𝒩 (0.5, 0.18) and clipped to the unit interval.

#### Human two-step task

To test whether the SCG reproduces canonical stay-probability signatures in the human two-step task, we simulated agents interacting with the task structure described above^49^ and analyzed behavior using conditional stay probabilities as a function of previous trial outcome and transition type.

We modeled a first-stage action *a* as deterministically selecting a distinct first-stage decision state *s*_*a*_. State values were computed from the current sparse cognitive graph, and first-stage action values were defined as

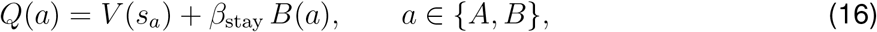

where *s*_*a*_ denotes the state associated with action *a, V* (*s*_*a*_) is the value of that state computed from the SCG, and *B*(*a*) = 1 if action *a* was chosen on the previous trial and 0 otherwise, capturing a standard stay bias.^49,62^ Choices followed a softmax policy,

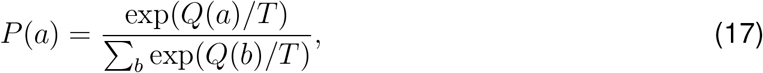

with temperature *T* controlling choice stochasticity. Second-stage choices were modeled as ideal selections of the option with the higher reward probability, while reward vectors, including those at second-stage states (*C* or *D*), were learned via a standard delta rule. On each trial, the agent chose between *A* and *B* using the softmax policy described above and updated *W, G*, and *r* according to the SCG learning rules at the experienced transitions.

We conducted a grid search over parameters targeting the conditional stay probabilities (transition × outcome) reported in.^49^ Results presented in **Figure 3** were generated from simulations using a representative parameter set that reproduced the canonical stay-probability interaction with *α*_→*R*_ = 0.6, *α*_→*NoR*_ = 0.3, *α*_*r*_ = 0.9, *γ* = 0.6, *T* = 0.5, *ζ* = 0.6, and *β*_*stay*_ = 0.6.

#### Mouse two-step task and optogenetic stimulation

We applied the SCG model to the mouse two-step task dataset reported in,^50^ using a hierarchical Bayesian modeling framework. The model state space was defined to reflect the animals’ experienced task structure:

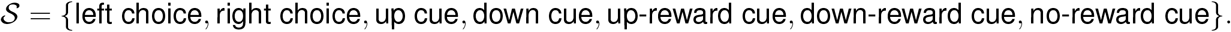

Choices (left vs. right) were modeled using the same action–state mapping (Eq. (16)) and choice policy (Eq. (17)) as in the human two-step task. Following prior analyses of this dataset^50^ and related modeling work,^51,62^ we assumed a *forgetful* reward representation, in which all elements of the reward vector were updated at a common rate, regardless of whether the corresponding state was visited. A non-forgetting variant of the SCG model was also evaluated but performed worse in terms of model evidence assessed by iBIC.

To estimate conditional stay probabilities (**Fig. 4C**), we simulated the SCG with fitted parameters under the same experimental conditions as the behavioral data. Transition learning-rate estimates shown in **Fig. 4D** were taken directly from session-by-session MAP estimates.

We modeled optogenetic stimulation in ChR2 animals at outcome timing as a boost in transition learning rates. We allowed distinct transition learning rates on stimulated trials, *α*_*s*,→*R*_ and *α*_*s*,→*NoR*_, in addition to the baseline learning rates *α*_→*R*_ and *α*_→*NoR*_ governing non-stimulated trials. Model predictions shown in **Fig. 4G** were generated using a representative parameter set (*α*_→*R*_ = 0.2, *α*_→*NoR*_ = 0.1, *α*_*r*_ = 0.5, *γ* = 0.4, *T* = 0.3, *ζ* = 0.6, *β*_stay_ = 0.4, *α*_*s*,→*R*_ = 0.7, *α*_*s*,→*NoR*_ = 0.3), chosen to illustrate the qualitative effects predicted by the model.

For comparison, we additionally fit several alternative models to the mouse data, including: (i) an SCG variant with a single transition learning rate (*α*_→*R*_ = *α*_→*NoR*_), (ii) an SCG variant without reward forgetting, (iii) the winning reinforcement-learning model reported in the original study (model-based reinforcement learning with asymmetric reward learning and forgetting),^50^ as well as standard successor representation, model-free, and model-based reinforcement-learning agents. Comparative results are shown in **Fig. S4**.

#### Hierarchical Bayesian model fitting

To robustly estimate model parameters ***h***, we performed a hierarchical Bayesian random-effects analysis^28,62^ separately for each animal. In this framework, the (appropriately transformed) parameter vector ***h***_*i*_ for experimental session *i* is treated as a random draw from a Gaussian population distribution with mean and covariance ***θ*** = {***µ***_*θ*_, **Σ**_*θ*_}. The subject-level parameter ***θ*** is estimated by maximizing the marginal likelihood

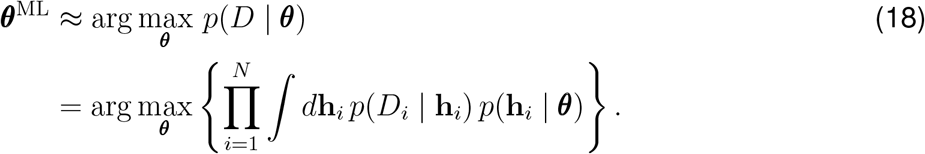

We optimized ***θ*** using an approximate Expectation–Maximization (EM) procedure. In the E-step of iteration *k*, a Laplace approximation yields

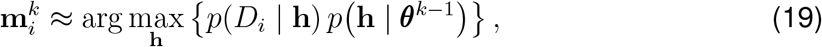

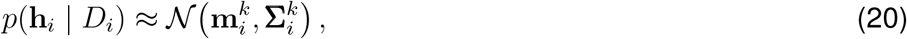

where 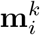 is the maximum-a-posteriori (MAP) estimate and 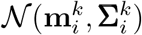 denotes a multivariate normal distribution with mean 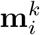 and covariance 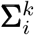, obtained from the inverse Hessian evaluated at 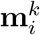.

In the M-step, population parameters are updated as

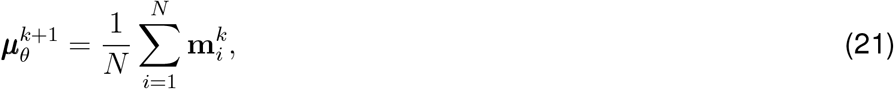

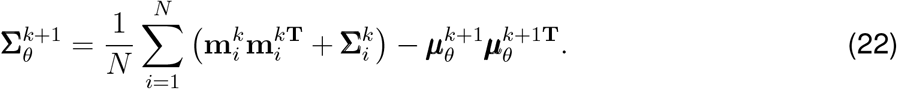

For simplicity, we assumed a diagonal covariance matrix **Σ**_*θ*_, corresponding to independent population-level effects.

When fitting the model to mouse data,^50^ model parameters were estimated by maximizing the log-likelihood over all free-choice trials.

#### Model comparison

Models were compared using the integrated Bayesian Information Criterion (iBIC).^28,62^ The marginal log likelihood of model *M* is given by

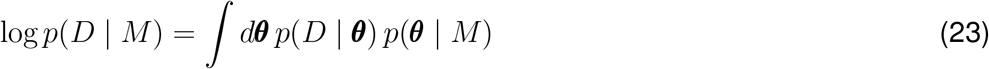

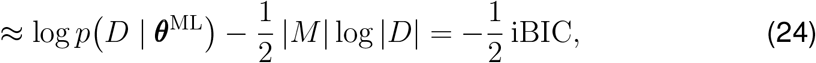

where |*M*| denotes the number of fitted population-level parameters and |*D*| is the total number of observed choices across all subjects.

The marginal likelihood term log *p*(*D*|***θ***^ML^) was computed by integrating out individual-level parameters:

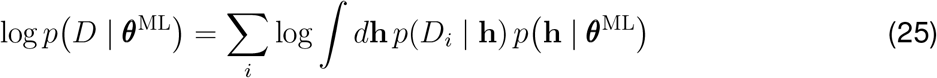

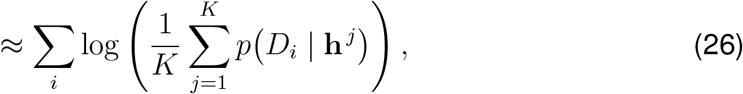

where the integral was approximated by Monte Carlo averaging over *K* samples **h** ^*j*^ drawn from the fitted prior *p*(**h** | ***θ***^ML^).

### M.5 Behavioral data analyses of the impact of optogenetic stimulation^50^

We focused on sessions in which optogenetic stimulation was delivered at the time of outcome onset and restricted analyses to balanced reward blocks (50%/50%) to minimize the influence of blockwise reward biases. For each free-choice trial *t*, we computed the probability of repeating the same first-stage choice on the next free-choice trial *t* + 1, *P* (stay_*t*+1_), conditioned on second-step transition (common vs. rare), outcome (reward vs. no reward), and stimulation (on vs. off). Because stimulation was experimentally constrained to be followed by at least two non-stimulated trials, these intervening trials were excluded; analyses were restricted to free-choice *t* and *t* + 1.

For descriptive comparisons, we contrasted *P* (stay_*t*+1_) following rare–reward trials with vs. without stimulation. Statistical significance of stimulation effects was assessed using permutation tests implemented in Python (SciPy). To quantify the joint influence of group and trial events, we fit a mixed-effects logistic regression,

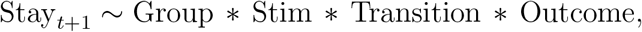

where Group (ChR2 vs. YFP), Stim (on vs. off), Transition (common vs. rare), and Outcome (reward vs. no reward) were entered as fixed effects, and mouse identity was modeled as a random effect. Analyses were restricted to free-choice trials, and permutation-based tests were specifically applied to compare stimulated vs. non-stimulated rare–reward trials.

### M.6 Spectral analysis of SCG

To characterize low-dimensional structure in the learned SCG, we performed spectral analyses of the directed adjacency matrix *G*. Because *G* is generally non-symmetric, we analyzed both right eigenvectors, defined by *Gv* = *λv*, and left eigenvectors, defined by *u*^⊤^*G* = *λu*^⊤^ (equivalently, eigenvectors of *G*^⊤^). Right eigenvectors emphasize structure associated with outgoing connectivity, whereas left eigenvectors emphasize structure associated with incoming connectivity. Eigenvectors associated with zero eigenvalues correspond to the null spaces of *G* and *G*^⊤^. These null spaces identify directions in state space that are not propagated under repeated application of the adjacency operator. In acyclic or sparsely connected graphs, such modes are typically associated with states lacking outgoing (terminal) or incoming (initial) connectivity, reflecting boundary structure or net-zero flow in the learned graph.

Spectral modes were examined for localization and periodicity across states under different task conditions. In particular, we compared eigenstructure across settings that differed in graph sparsity and cyclicity, including cases in which transitions through an inter-trial interval (ITI) state were weak or absent, and cases in which training induced recurrent transitions linking reward states to subsequent trial onset.

For reference, we also performed the same analyses on the dense transition representation *W*. In contrast to the sparsified adjacency matrix *G*, the matrix *W* encodes graded long-horizon transition statistics accumulated across multiple paths. As a result, its spectral structure reflects averaged predictive flow rather than discrete reachability boundaries. In directed but non-cyclic task configurations, eigenmodes of *W* can exhibit partial localization at source or sink states. This arises because, in absorbing or strongly directed systems, the dominant eigenstructure of a predictive operator is shaped by imbalance between incoming and outgoing predictive mass, leading to accumulation near terminal states. However, because *W* retains contributions from many indirect transitions, these modes are typically smoother and less sharply localized than those obtained from the thresholded graph *G*.

When task dynamics induce approximate translational invariance through recurrent transitions, the spectrum of *W* is dominated by low-frequency modes associated with repeated structure, yielding periodic eigenmodes analogous to those previously described for the successor representation in spatial random walk settings.^3^ In this regime, spectral modes primarily reflect global predictive geometry rather than discrete start or terminal roles. By contrast, sparsification into *G* suppresses weak and indirect transitions, accentuating boundary structure and emphasizing directed reachability relevant for planning and valuation.

All spectral analyses were performed directly on adjacency matrices rather than graph Laplacians,^3^ which are more appropriate for undirected or diffusion-based analyses, as the learned representations were explicitly directed. This ensured that extracted eigenmodes reflected the directed reachability structure underlying computation in the SCG.

### M.7 Simulations of grid- and flag-like representations

To visualize boundary-localized eigenmodes in a cognitive task, we used the learned *G* from the mouse two-step environment with seven states and threshold *ζ* = 0.6. We computed left and right eigenvectors of *G* and visualized on a schematic representation of the task, using Gaussian interpolation for display purposes.

To examine conditions under which periodic structure can emerge in a non-spatial task, we extended the state space with an inter-trial interval (ITI) state that returned to left or right with equal probability, *P* (left|ITI) = *P* (right|ITI) = 0.5, and used lower threshold *ζ* = 0.5. This produced a cyclic graph spanning trial onset, transition, outcome, ITI, and subsequent trial onset. Spectral decomposition of *G* in this augmented graph yielded periodic eigenmodes along the trial sequence, analogous to grid-like structure in abstract state spaces.

At higher sparsification thresholds, the adjacency matrix *G* no longer preserved cyclic structure to support periodic eigenmodes. In these cases, however, periodic structure remained evident in spectral decompositions of the dense transition representation *W*, reflecting its sensitivity to translational symmetry in the underlying transition statistics even when such structure is suppressed by graph sparsification.

For comparison, we simulated a 12 × 12 spatial grid world in which agents moved uniformly in up to four directions with reflective boundaries.

## Author Contributions

AG, PS, and KI conceived the project and developed the model, with input from IA. AG, PS, IA, and KI performed model analyses and interpreted the results. PS led empirical data analysis and model fitting, with assistance and input from AG, IA, MBP, KI. MBP collected the mouse data in the original study and restructured the dataset for the present analyses. AG, PS, IA, MBP and KI discussed the results and wrote the manuscript.

## Acknowledgments

We thank Mark Walton and Thomas Akam for sharing the mouse dataset and for valuable input on the analysis. We thank Evan Russek, Ida Momennejad, Nathaniel Daw, and Sam Gershman for sharing human structural learning datasets. We thank Peter Balsam, C. Randy Gallistel, Kim Stachenfeld, Peter Dayan, Matt Schaefer, and Deniz Urey for valuable suggestions. Research reported in this publication was supported by the National Institute of Mental Health (R01MH136214; AG, PS, KI), the Brain & Behavior Research Foundation Young Investigator Grant (KI), and the National Institute of Neurological Disorders and Stroke (T32NS064929; IA). Mouse data collection was supported by a Wellcome Trust studentship to MBP (215198/Z/19/Z) and a Wellcome Trust Senior Research Fellowship to Mark Walton (202831/Z/16/Z).

## Supplementary Figures

**Figure S1:**
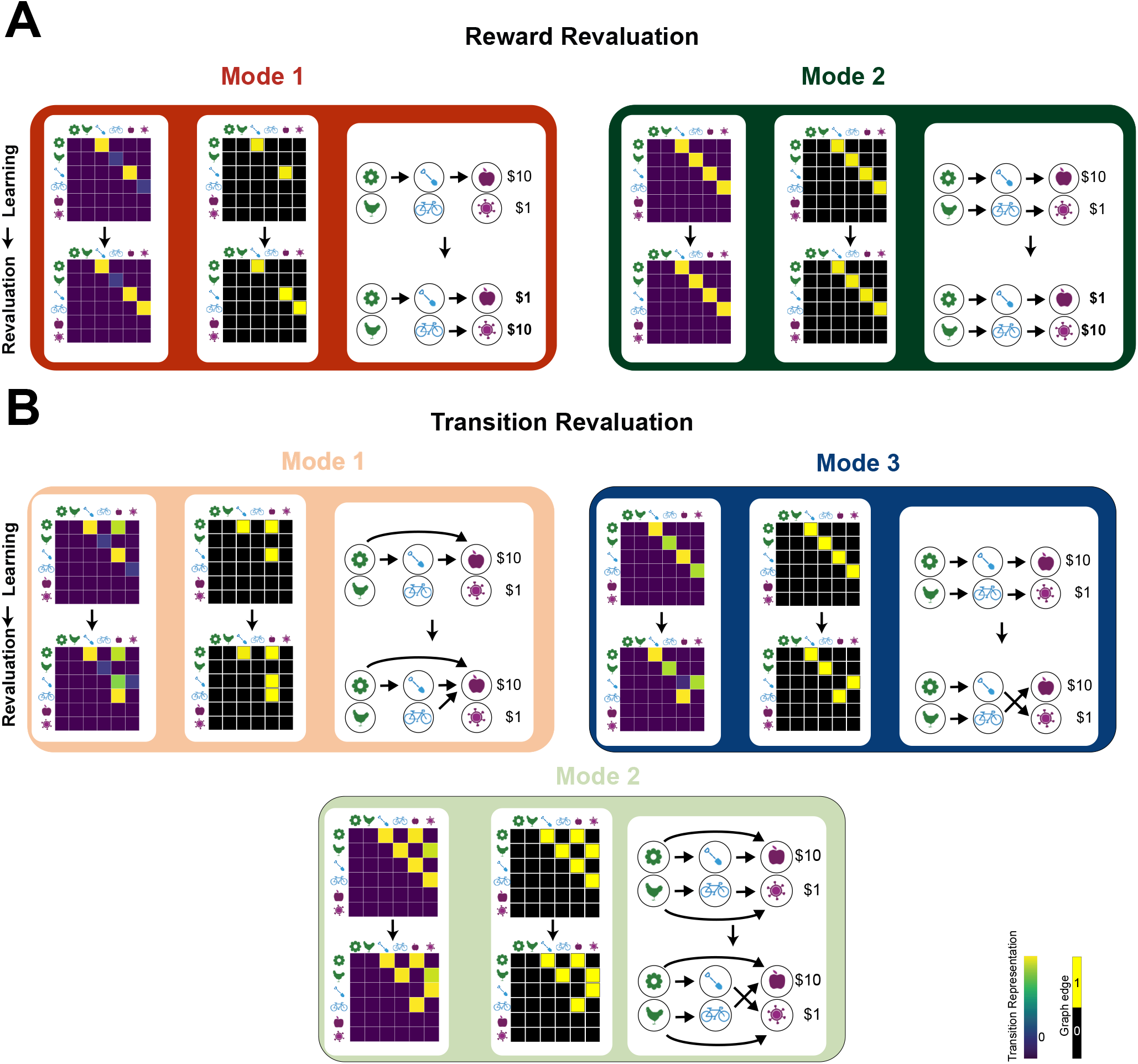
Transition representation *W* and thresholded adjacency matrix *G* that underlie multimodal distributions in **(A)** reward revaluation and **(B)** transition revaluation.

**Figure S2:**
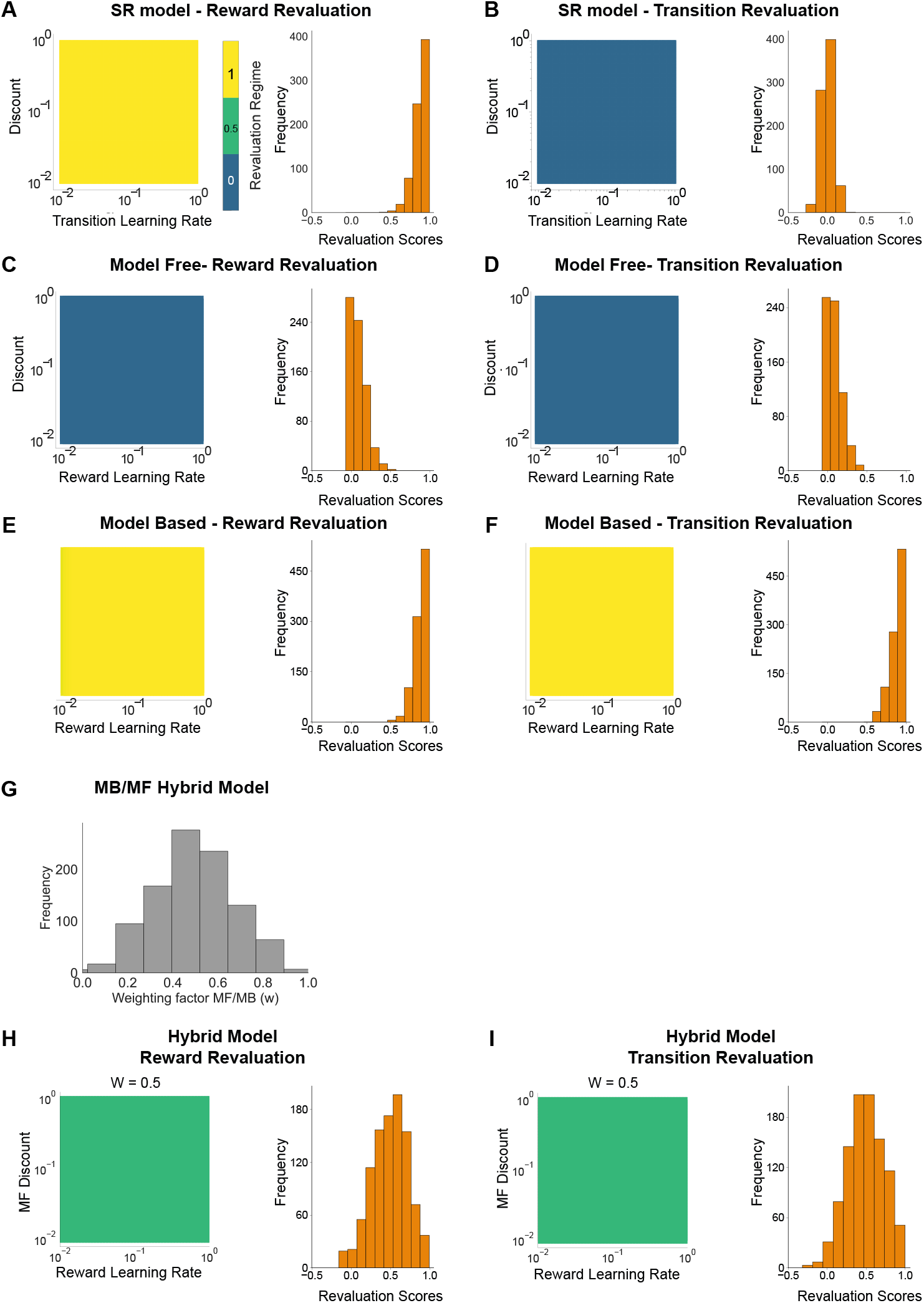
Alternative models fail to capture multimodal behavioral regimes in reward and transition revaluation tasks.^48^ (**A**) and (**B**). Successor representation (SR) model. Phase diagrams (left) and predicted revaluation score distributions (right). In the reward revaluation task, the SR model predicts a uniform change in preference for the initial states across parameter regimes (**A**, left), yielding a unimodal distribution of revaluation scores centered at 1 (complete change; **A**, right). In contrast, in the transition revaluation task, the SR model predicts no change in preference for the initial states regardless of parameters (**B**, left), resulting in a unimodal distribution centered at 0 (no change; **B**, right). (**C**) and (**D**). Model-free reinforcement learning (temporal-difference learning). The model predicts no change in preference for both reward and transition revaluation tasks across parameter regimes. (**E**) and (**F**). Model-based reinforcement learning. A transition model combined with forward planning predicts complete reversal of preferences in both tasks. (**G**). Hybrid model-based/model-free (MB/MF) models with a distributed weighting parameter between model-based and model-free controls (left) fail to reproduce the multimodal distributions observed in human behavior for either reward or transition revaluation (right).

**Figure S3:**
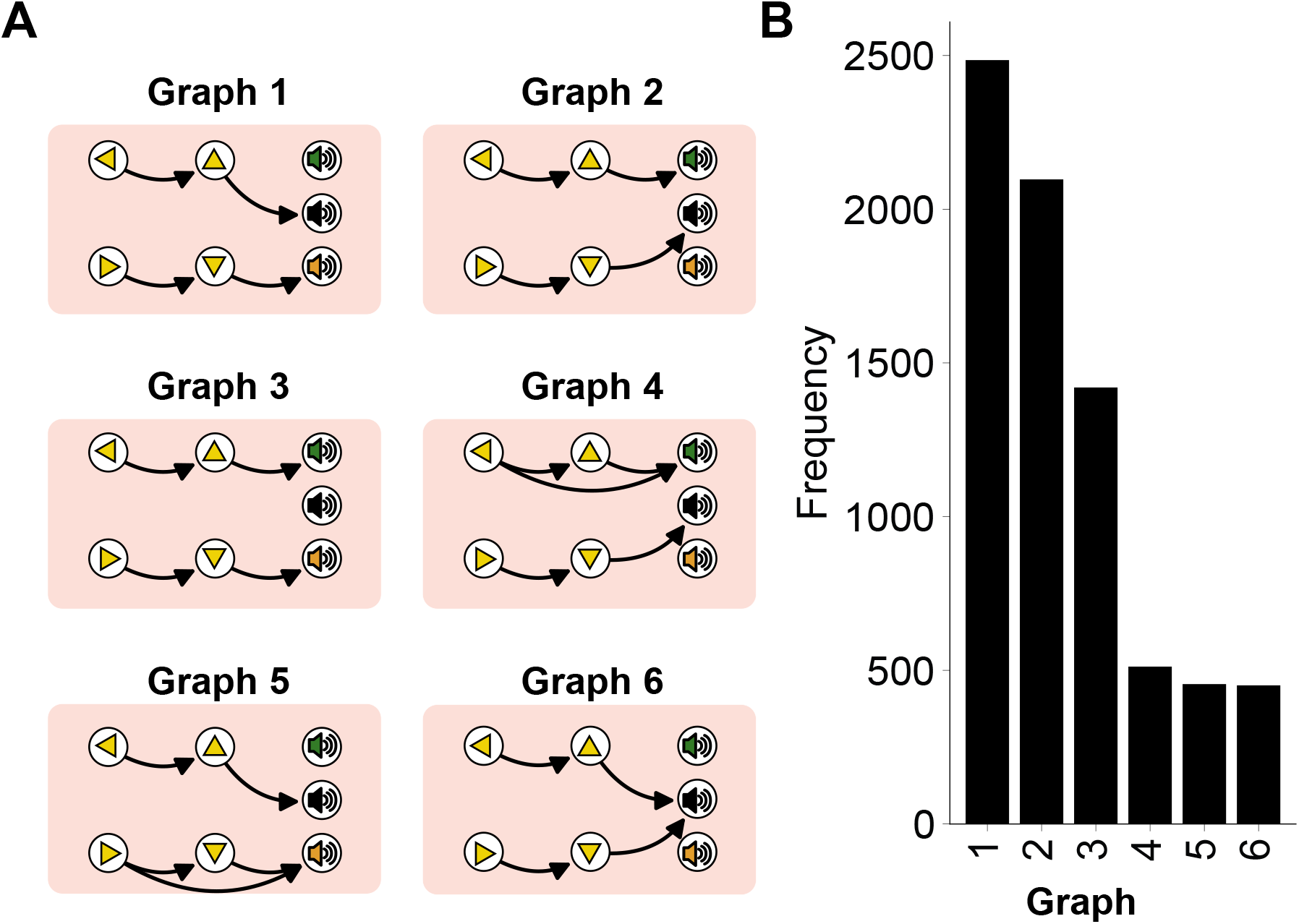
Graph configuration frequencies in the mouse two-step task. (**A**). Most frequently occurring graph configurations. (**B**). Histogram of graph configuration frequencies for the representative animal.

**Figure S4:**
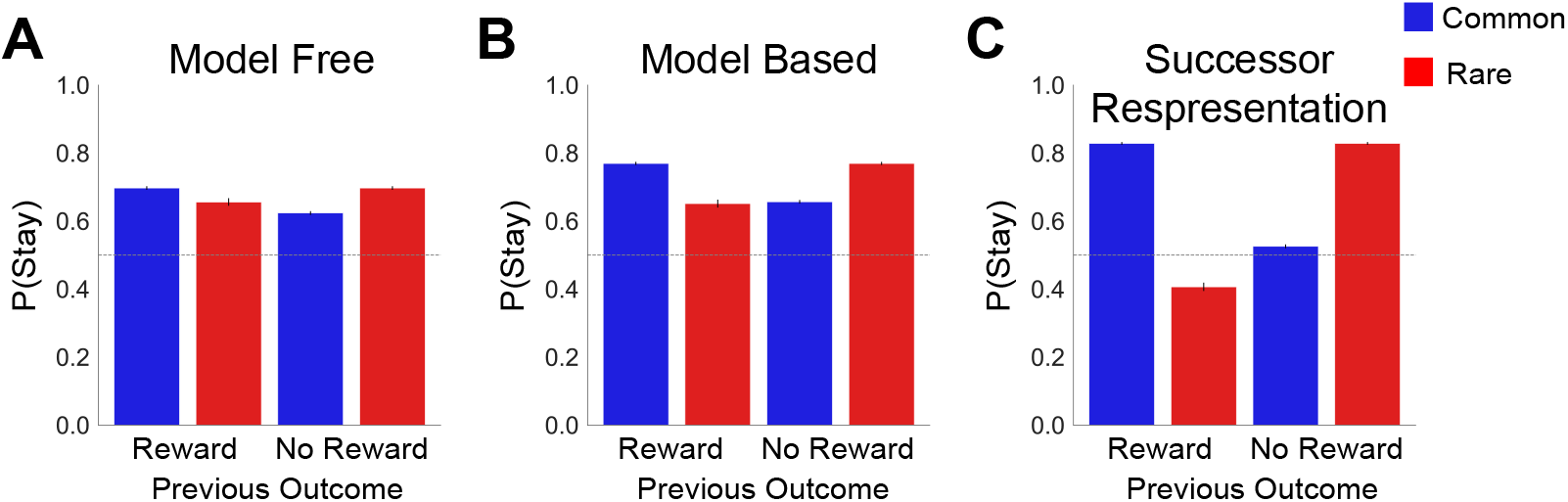
Alternative models fail to reproduce mouse two-step behavior. (**A**) Standard model-free temporal difference learning model. (**B**) Standard model-based reinforcement learning model. (**C**) Standard SR model. Error bars indicate s.e.m. across simulations.

**Figure S5:**
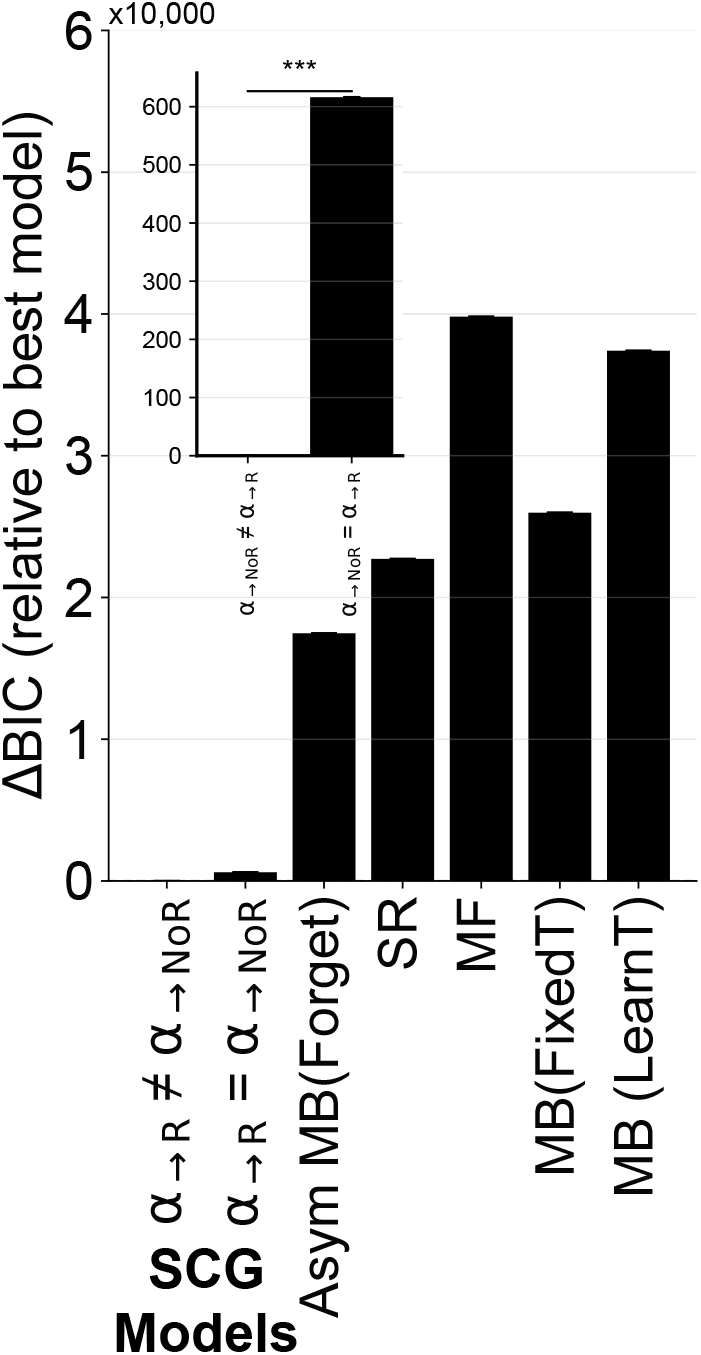
Model comparison for mouse two-step task.^50^ Integrated Bayesian Information Criterion (iBIC) scores are shown for models fit using a hierarchical Bayesian approach. The SCG model with separate transition learning rates for rewarded and unrewarded outcomes (*α*_→R_ and *α*_→NoR_) outperformed the SCG model with a single transition learning rate (*α*_→R_ = *α*_→NoR_), as well as the winning model of previous analysis (model-based RL with asymmetric learning and forgetting^50^) standard SR, standard model-free, and standard model-based agents with fixed transition structure and learned transition structure. The inset shows a zoomed view of the iBIC differences between SCG variants. Lower iBIC values indicate better model fit. Error bars indicate the cross-sample mean ± s.e.m. (4,000 bootstrap samples from the posterior distribution)

**Figure S6:**
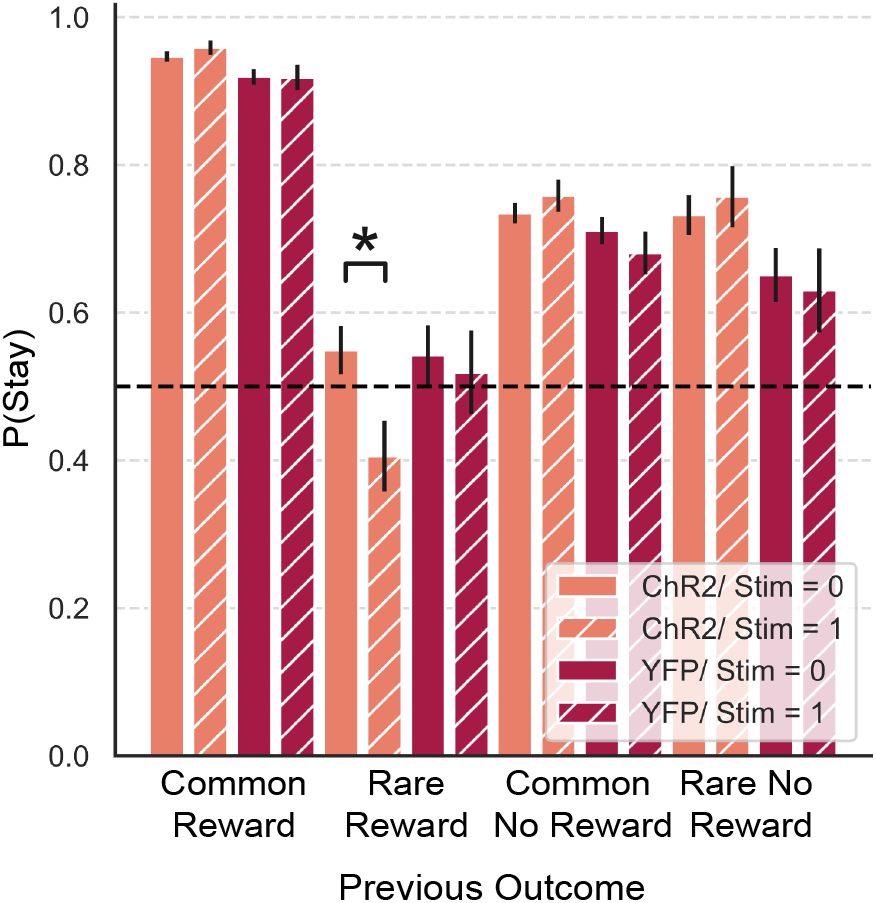
Optogenetic stimulation effects on conditional stay probability.^50^ Optogenetic stimulation at outcome timing in the mouse two-step task is associated with differences in subsequent choice behavior following rare transitions. Choice probabilities are shown for ChR2 (orange) and YFP control (red) groups, comparing trials with stimulation (shaded bars) and without stimulation (solid bars) across transition and outcome conditions. Error bars indicate s.e.m..

**Figure S7:**
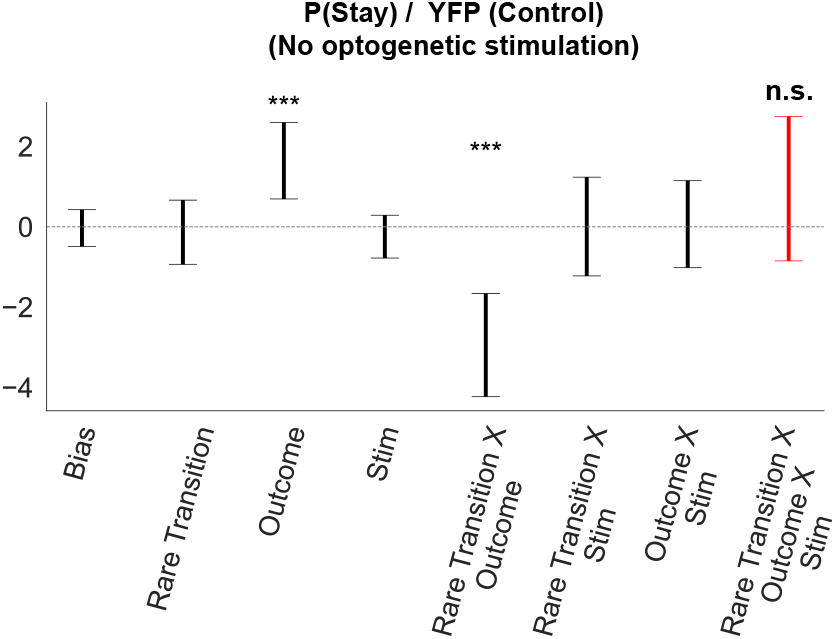
Regression analysis of dopamine stimulation effects in control animals. Unlike the ChR2 group, the three-way interaction (red), associated with enhanced transition learning from stimulation, did not differ significantly from chance levels in control animals (*p* = 0.3). Error bars show ±2 SE (approx. 95 % CI).

**Figure S8:**
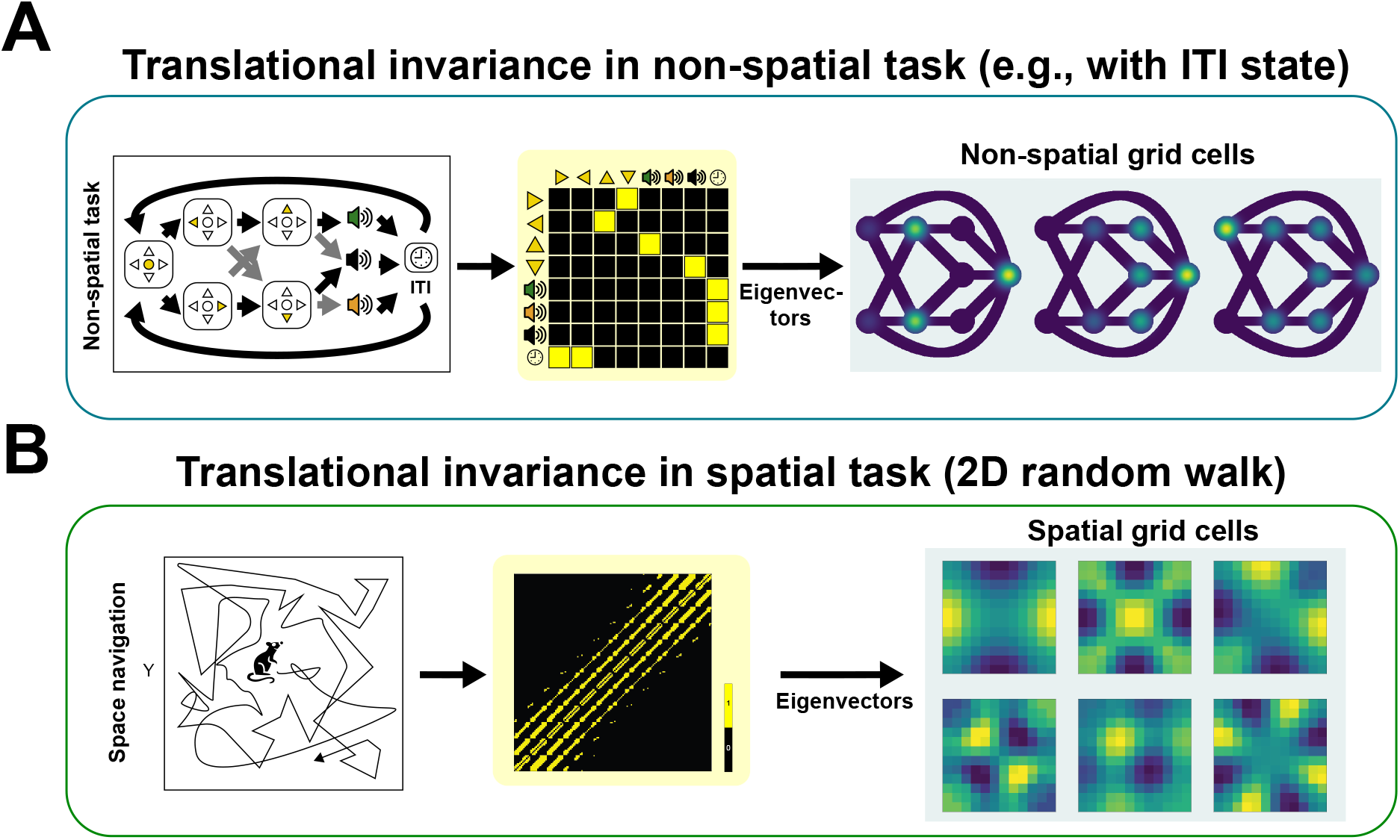
Grid-like and flag-like neural codes predicted by the SCG. (**A**) Grid-like codes in non-spatial tasks. Grid-like patterns emerge in trial-based decision tasks with temporal translational invariance. When the SCG forms cyclic graph structure, for example by including the inter-trial interval, it predicts grid-like activity over an abstract task space. (**B**) Grid-like codes in spatial tasks can also appear from the SCG with a relatively low threshold value.

